# The interferon stimulated gene-encoded protein HELZ2 inhibits human LINE-1 retrotransposition and LINE-1 RNA-mediated type I interferon induction

**DOI:** 10.1101/2022.03.26.485892

**Authors:** Ahmad Luqman-Fatah, Yuzo Watanabe, Fuyuki Ishikawa, John V. Moran, Tomoichiro Miyoshi

## Abstract

Some interferon stimulated genes (ISGs) encode proteins that inhibit LINE-1 (L1) retrotransposition. Here, we used immunoprecipitation followed by liquid chromatography-tandem mass spectrometry to identify proteins that associate with the L1 ORF1-encoded protein (ORF1p) in ribonucleoprotein particles. Three ISG proteins that interact with ORF1p inhibit retrotransposition: HECT and RLD domain containing E3 ubiquitin-protein ligase 5 (HERC5); 2’-5’-oligoadenylate synthetase-like (OASL); and helicase with zinc finger 2 (HELZ2). HERC5 destabilizes ORF1p, but does not affect its cellular localization. OASL impairs ORF1p cytoplasmic foci formation. HELZ2 recognizes sequences and/or structures within the L1 5’UTR to reduce L1 RNA, ORF1p, and ORF1p cytoplasmic foci levels. Overexpression of WT or reverse transcriptase-deficient L1s led to a modest induction of IFN-α expression, which was abrogated upon HELZ2 overexpression. Notably, IFN-α expression was enhanced upon overexpression of an ORF1p RNA binding mutant, suggesting ORF1p binding might protect L1 RNA from “triggering” IFN-α induction. Thus, ISG proteins can inhibit retrotransposition by different mechanisms.

## Introduction

Sequences derived from Long INterspersed Element-1 (LINE-1 or L1) retrotransposons comprise ~17% of human genomic DNA^1^. The overwhelming majority of L1-derived sequences have been rendered retrotransposition-defective by mutational processes either during or after their integration into the genome^2–4^. However, an average human genome is estimated to contain at least 100 full-length human-specific retrotransposition-competent L1s (RC-L1s)^5–7^, with only a small number of human-specific “hot” L1s responsible for the bulk of retrotransposition activity^6,8^.

Human RC-L1s are ~6 kb and consist of a 5’ untranslated region (UTR), two open reading frames (ORF1 and ORF2), and a 3’UTR that ends in a poly(A) tract^4,9,10^. ORF1 encodes a ~40 kDa protein (ORF1p) that has RNA-binding and nucleic acid chaperone activities^11–13^. ORF2 encodes a ~150-kDa protein (ORF2p) that has endonuclease (EN) and reverse transcriptase (RT) activities required for canonical L1 retrotransposition^14–17^. RC-L1s mobilize via a “copy-and-paste” mechanism, where an L1 RNA intermediate is reverse transcribed into an L1 cDNA at a new genomic integration site by a process termed target-site primed reverse transcription (TPRT)^16,18–20^.

L1 retrotransposition begins with transcription of full-length RC-L1 sense strand RNA using an internal RNA polymerase II promoter located within the L1 5’UTR^21–23^. The resultant bicistronic L1 mRNA is exported to the cytoplasm, where its translation leads to the production of ORF1p and ORF2p. ORF1p and ORF2p preferentially associate with their encoding L1 RNA, by a process known as *cis*-preference^24,25^, to form a cytoplasmic L1 ribonucleoprotein (RNP) complex that appears necessary, but not sufficient for retrotransposition^26,27^. Components of the L1 RNP gain access to the nucleus by a process that does not strictly require mitotic nuclear envelope breakdown^28^, although recent reports suggest that components of the L1 RNP might also gain access to genomic DNA during mitotic nuclear envelope breakdown^29^.

Once in the nucleus, ORF2p EN makes a single-strand endonucleolytic nick at a consensus target sequence (*e.g.*, 5’-TTTTT/AA-3’ and related variants of that sequence) in genomic DNA, generating 5’-PO_4_ and 3’-OH groups^16,17,20,30,31^. Base pairing between the short stretch of thymidines in genomic DNA liberated by L1 EN cleavage and the 3’ L1 poly(A) tract is thought to form a primer/template complex^27,32^, where the 3’-OH group of genomic DNA serves as a primer to allow ORF2p RT to generate (−) strand L1 cDNA from its associated L1 RNA template^16,17,19,32^. How top strand genomic DNA cleavage and (+) strand L1 cDNA synthesis occurs requires elucidation, but each step likely requires activities contained within ORF2p^4,33–35^. The completion of TPRT results in the integration of an L1 at a new genomic location.

L1 retrotransposition is mutagenic and, on rare occasions, can lead to human genetic diseases^4,36–39^. Besides acting as an insertional mutagen, products generated during the process of L1 retrotransposition (*e.g.*, double-stranded L1 RNAs and single-stranded L1 cDNAs) are hypothesized to trigger a type I interferon (IFN) response that may contribute to inflammation and aging phenotypes^40–46^. However, how L1 expression contributes to the induction of a type I IFN response and whether this process plays a direct role in human diseases require elucidation.

Previous studies revealed that ORF1p, ORF2p, and L1 RNA can localize within cytoplasmic foci that often are in close proximity to stress granules (SGs) – dynamic membraneless cytoplasmic structures that form upon the treatment of cells with certain stressors – although it is unclear what role, if any, cytoplasmic foci play in L1 biology^47–50^. SGs sequester polysomes, host proteins, and cellular RNAs and are proposed to function as regulatory hubs during the cellular stress response^51,52^. Intriguingly, host factors that inhibit L1 retrotransposition (*e.g.*, the zinc-finger antiviral protein [ZAP] or MOV10 RNA helicase) frequently co-localize with L1 cytoplasmic foci^50,53,54^.

To further understand the suite of host factors that bind to L1 RNPs, we generated a panel of ORF1p missense mutation and tested them for their ability to: (1) be stably expressed in human cell lines; (2) reduce the formation of cytoplasmic foci; (3) impair the ability to bind L1 RNA; and (4) inhibit L1 retrotransposition. These analyses led to the identification of a triple mutant, R206A/R210A/R222A (a.k.a., M8/RBM), in the ORF1p RNA binding domain^12^.

Immunoprecipitation (IP) coupled with liquid chromatography-tandem mass spectrometry (LC-MS/MS) analyses followed by Gene Ontology (GO)^55^ and Gene Set Enrichment Analysis (GSEA)^56^ revealed that a full-length RC-L1 containing a carboxyl-terminal epitope-tagged version of ORF1p (WT ORF1p-FLAG) preferentially associates with proteins encoded by several interferon stimulated genes (ISGs), including HERC5, HELZ2, OASL, DDX60L, and IFIT1. Detailed analyses revealed that HERC5, HELZ2, and OASL overexpression inhibits the retrotransposition of engineered L1s in cultured cells and that each protein appears to act at different steps in the L1 retrotransposition cycle. Finally, we report that HELZ2 preferentially recognizes RNA sequences and/or RNA structures within the L1 5’UTR to destabilize L1 RNA and that HELZ2 overexpression reduced the abilty of engineered L1 RNAs to induce IFN-α expression.

## Results

### Construction of a panel of ORF1p missense mutations

To refine the role of ORF1p domains necessary for L1 retrotransposition and/or cytoplasmic foci formation, we generated a panel of ORF1p alanine missense mutations in a full-length human RC-L1 expression construct that expresses a version of ORF1p containing a FLAG epitope tag at its carboxyl-terminus (**Fig. 1a**, pJM101/L1.3FLAG)^5,50^. Mutations were generated in the following ORF1p regions: (1) M1: a conserved pair of amino acids (N157A/R159A) important for ORF1p cytoplasmic foci formation and L1 retrotransposition^48^; (2) M2: a pair of amino acids predicted to play a role in ORF1p trimerization^57^ (R117A/E122A); (3) M3 and M4: amino acids proposed to mediate the coordination of chloride ions in the coiled-coil domain to stabilize ORF1p homotrimer formation^12^ (N142A and R135A, respectively); (4) M5: a putative ORF1p protein-protein interaction surface that may interact with host factors through its acidic patch^12^ (E116A/D123A); (5) M6-M9: amino acids required for ORF1p RNA binding activity^12,26,49^ (K137A/K140A, R235A, R206A/R210A/R211A, and R261A, respectively); and (6) M10: an amino acid thought to decrease nucleic acid chaperone activity^49^ (Y282A). The relative position of each mutation in the ORF1p crystal structure^12^ and the putative functions of the wild type amino acids are shown in **Supplementary Figs. 1 and 2a**.

**Figure. 1:**
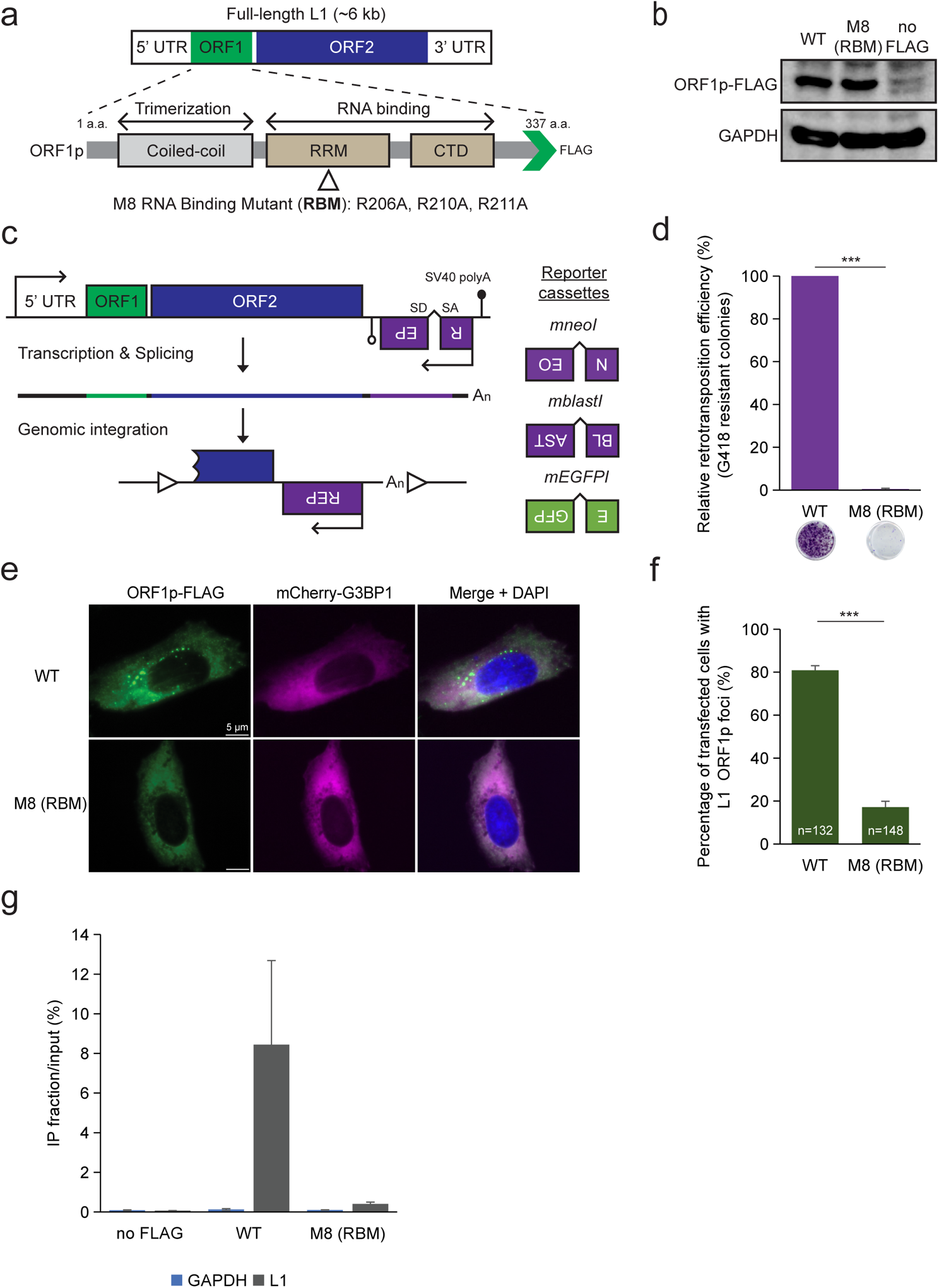
Identification of an ORF1p RNA-binding mutant critical for L1 retrotransposition and ORF1p cytoplasmic foci formation. **(a)** *Schematic of a full-length RC-L1 (L1.3: Genbank Accession #L19088).* ORF1p functional domains are noted below the schematic and include the coiled-coil domain, the RNA recognition motif (RRM), and carboxyl-terminal domain (CTD). Green arrowhead, position of the in-frame FLAG epitope tag. Open triangle, relative position of a triple mutant (R206A/R210A/R211A) in the RRM domain. **(b)** *WT ORF1p and the ORF1p-FLAG R206A/R210A/R211A mutant are stably expressed in HeLa-JVM cells*. Western blot with an anti-FLAG antibody. A construct lacking the FLAG epitope tag (pJM101/L1.3 [no FLAG]) served as a negative control. GAPDH served as a loading control. **(c)** *Schematics of the retrotransposition indicator cassettes used in this study.* A retrotransposition indicator cassette (*REP*) was inserted into the 3’UTR of an L1 in the opposite orientation relative to sense strand L1 transcription. The *REP* gene contains its own promoter (upside down arrow) and polyadenylation signal (open lollipop). The *REP* gene is interrupted by intron in the same orientation relative to sense strand L1 transcription. This arrangement ensures that *REP* expression only will occur if the sense strand L1 transcript is spliced and successfully integrated into genomic DNA by retrotransposition (bottom schematic, open triangles, target site duplications that typically are generated upon L1 retrotransposition). Three retrotransposition indicator cassettes are shown at the right of the figure: *mneoI*, which confers resistance to G418; *mblastI*, which confers resistance to blasticidin; and *mEGFPI*, which leads to enhanced green fluorescent protein (EGFP) expression. **(d)** *Results of a representative mneoI-based retrotransposition assay*. HeLa-JVM cells were co-transfected with phrGFP-C (transfection control) and either pJM101/L1.3FLAG (WT) or pALAF008 (M8 [RBM]). X-axis, L1 construct names and representative retrotransposition assay results. Y-axis, relative retrotransposition efficiency; the number of G418 resistant (retrotransposition-positive) foci was normalized to the transfection efficiency (*i.e.*, the percentage of hrGFP-positive cells). Pairwise comparison relative to the WT control: *p* = 2.1 × 10^−12^***. **(e)** *The ORF1p-FLAG R206A/R210A/R211A mutant (M8 [RBM]) reduces the number of ORF1p cytoplasmic foci.* Representative immunofluorescence microscopy images of U-2 OS cells expressing either WT ORF1p-FLAG (pJM101/L1.3-FLAG) or ORF1p-FLAG R206A/R210A/R211A mutant (pALAF008 [M8 (RBM)]). The U-2 OS cells also expressed a doxycycline-inducible (Tet-On) mCherry-G3BP1 protein. White scale bars, 5 µm. **(f)** *Quantification of immunofluorescence assays in U-2 OS* cells. X-axis, L1 construct names. Y-axis, percentage of transfected cells containing ORF1p cytoplasmic foci. The number (n) inside the green bars indicates the number of individual cells counted in the assay. Pairwise comparisons relative to the WT control: *p* = 7.5 × 10^−11^***. **(g)** *RNA-immunoprecipitation (RNA-IP) reveals an L1 RNA binding defect in the ORF1p-FLAG R206A/R210A/R211A mutant (M8 [RBM]).* HeLa-JVM cells were transfected with either pJM101/L1.3 (no FLAG), WT ORF1p-FLAG (pJM101/L1.3-FLAG), or the ORF1p-FLAG R206A/R210A/R211A mutant (pALAF008 [M8 (RBM)]). An anti-FLAG antibody was used to immunoprecipitate ORF1p-FLAG; reverse transcription-quantitative PCR (RT-qPCR) using a primer set (L1 [SV40]) that amplifies RNAs derived from the transfected L1 plasmid was used to quantify L1 RNA. X-axis, constructs name. Y-axis, the enrichment of L1 RNA levels between the IP and input fractions. Blue rectangles, relative levels of control GAPDH RNA (primer set: GAPDH). Gray rectangles, relative levels of L1 RNA. In panels **(d)**, **(f)**, and **(g)**, values represent the mean ± the standard error of the mean (SEM) of three independent biological replicates. The *p*-values were calculated using a one-way ANOVA followed by Bonferroni-Holm post-hoc tests; *** *p<*0.001.

### ORF1p RNA-binding is critical for ORF1p cytoplasmic foci formation

Western blot analyses, using an antibody that recognizes the ORF1p FLAG epitope tag, revealed that each of the ORF1p mutant constructs could be expressed in human U-2 OS osteosarcoma, HeLa-JVM cervical cancer, and HEK293T embryonic kidney cell lines (**Fig. 1b and Supplementary Fig. 2b**). We observed a severe reduction in the steady state level of ORF1p in the M1 mutant, as well as an alteration in the electrophoretic mobility of ORF1p in the M5 mutant, when compared to the WT ORF1p-FLAG control in each cell line (**Supplementary Fig. 2b**). The steady state levels of the M9 and M10 ORF1p mutant proteins appeared to be reduced in the HeLa-JVM and HEK293T cell lines, but not in the U-2 OS cell line, when compared to the WT ORF1p-FLAG control (**Supplementary Fig. 2b**).

We next assayed whether the ORF1p mutations affected L1 retrotransposition efficiency. Briefly, each of the full-length WT pJM101/L1.3FLAG and mutant ORF1p derivatives (mutants M1 to M10) constructs contain an *mneoI* retrotransposition indicator cassette within their 3’UTR, ensuring the G418-resistant foci will only arise upon the completion of a single round of retrotransposition^17^. The L1 retrotransposition efficiency was calculated by counting the resultant number of G418-resistant foci, which was normalized to the transfection efficiency, upon completion of the assays^17,58,59^ (**Figs. 1c and 1d; see Methods**).

The M1, M2, M5, M8, and M9 mutants exhibited severely reduced L1 retrotransposition efficiencies when compared to the positive control (*i.e.,* >90% the levels of pJM101/L1.3FLAG). By comparison, the M6, M7, and M10 mutants only exhibited a ~60 to 70% decrease in L1 retrotransposition efficiency, whereas the M3 and M4 mutants had no discernable effect on L1 retrotransposition efficiency, when compared to the pJM101/L1.3FLAG positive control (**Supplementary Fig. 2c**). A construct harboring a missense mutation within the ORF2p reverse transcriptase domain (D702A) served as a negative control. The above data suggest that the putative trimerization, RNA-binding, nucleic acid chaperone, and ORF1p protein-binding domains are important for L1 retrotransposition^11,12,17,25,49^. Because the M3 and M4 mutants did not show a reduction in L1 retrotransposition efficiency, these data suggest that single point mutations in the putative chloride-ion coordinating sites (R135A or N142A) are not sufficient to destabilize ORF1p trimerization when compared to either the M2 mutant or the G132I/R135I/N142I triple mutant used in a previous study^12^.

We next focused our analyses on the M2, M5, and M8 mutants because their respective versions of ORF1p are stably expressed in HeLa-JVM cells despite severely reducing L1 retrotransposition efficiency. To determine whether the M2, M5, and M8 mutant ORF1p proteins localize to cytoplasmic foci, we established a U-2 OS cell line that expresses a doxycycline-inducible stress granule protein, G3BP1, which is tagged at its amino terminus with a mCherry fluorescent protein (mCherry-G3BP1)^60^ (**Supplementary Fig. 3a**). The U-2 OS cells were transfected with either the WT (pJM101/L1.3FLAG), M2, M5, or M8 mutant ORF1p derivatives and ORF1p-FLAG was visualized ~48 hours post-transfection using an anti-FLAG primary antibody and Alexa Fluor 488-conjugated anti-mouse IgG secondary antibody (see Methods). The M2 and M5 mutants were able to form cytoplasmic foci at comparable numbers and intensities relative to the WT control (**Supplementary Fig. 3b**). An increase in size of the ORF1p cytoplasmic foci and the co-localization of ORF1p with the stress granule marker mCherry-G3BP1 was further enhanced upon arsenite treatment (**Supplementary Fig. 3c**). By comparison, the M8 ORF1p RNA binding mutant exhibited a severe reduction in the percentage of cells containing ORF1p cytoplasmic foci (~15% of cells) when compared to U-2 OS cells expressing either the WT, M2, or M5 constructs (~80% of cells) even though it was stably expressed in HeLa-JVM, U-2 OS, and HEK293T cells (**Figs. 1b, 1e, and 1f**; **Supplementary Figs. 2b, 3b, 3c, and 3d**). RNA-immunoprecipitation (RNA-IP) experiments confirmed that the M8 mutant was impaired for its ability to bind L1 RNA when compared to WT ORF1p (**Fig. 1g. and see below**), which is consistent with the previous study^12^.

To confirm that the M8 ORF1p protein exhibited reduced RNA binding, we transfected the pJM101/L1.3FLAG (ORF1p-FLAG) or pALAF008_L1.3FLAG_M8 (M8/RBM-FLAG) expression constructs into HeLa-JVM cells and immunoprecipitated (IP) the resultant ORF1p complexes using an anti-FLAG antibody (**Fig. 2a**). Control western blot experiments revealed a similar level of WT and M8/RBM ORF1p-FLAG in whole cell extracts and immunoprecipitates from the HeLa-JVM whole cell extracts cells, but not in a negative control transfected with an L1 expression vector lacking the FLAG epitope tag (**Fig. 2b**). Moreover, the Poly(A) Binding Protein Cytoplasmic 1 (PABPC1) was robustly detected in IP reactions conducted with WT ORF1p-FLAG cell extracts, but was severely reduced in IP reactions conducted with M8/RBM ORF1p-FLAG L1 cell extracts (**Fig. 2b**), which is consistent with previous studies that found the association between ORF1p and PABPC1 requires RNA^50,61^. Thus, the above data suggest that the M2, M5, and M8 mutants each produce similar steady state levels of ORF1p and reduce L1 retrotransposition efficiencies. However, cytoplasmic foci formation depends on the ability of ORF1p to bind RNA (M8 mutant). Given these data, we focused our subsequent studies on the WT ORF1p-FLAG and M8/RBM-FLAG proteins (herein called the RNA Binding Mutant [RBM]).

**Figure. 2:**
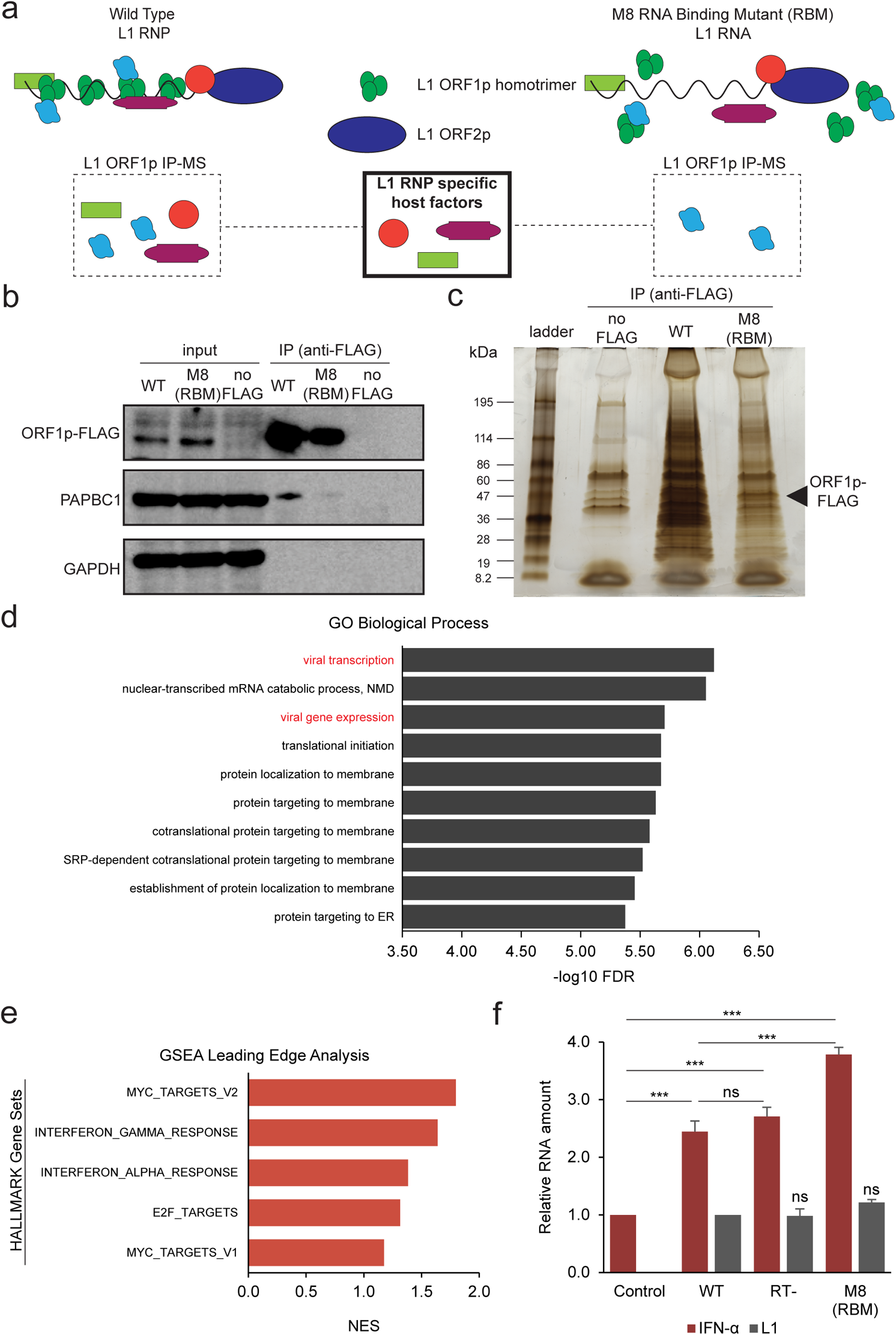
The proteins encoded by interferon-responsive genes are enriched in WT ORF1p-FLAG, but not ORF1p-FLAG (M8 [RBM]) mutant complexes. **(a)** *Experimental rationale for identifying host factors enriched in WT ORF1p-FLAG vs. ORF1p-FLAG (M8 [RBM]) immunoprecipitation reactions.* Hypothetical diagrams of the proteins associating with WT and M8 (RBM) mutant RNP particles. Green circles, ORF1p-FLAG. Blue Oval, ORF2p. Red circle, purple squared oval, and green rectangle, host factors that might associate with ORF1p-FLAG and/or L1 RNPs. **(b)** *The ORF1p (M8 [RBM]) mutant does not efficiently interact with Poly(A) Binding Protein Cytoplasmic 1 (PABPC1)*. HeLa-JVM cells were transfected with either pJM101/L1.3 (no FLAG), pJM101/L1.3-FLAG (WT ORF1p-FLAG), or pALAF008 (ORF1p-FLAG [M8 [RBM]] mutant). An anti-FLAG antibody was used to immunoprecipitate ORF1p-FLAG. Western blots detected ORF1p (anti-FLAG), PABPC1 (anti-PABC1), and GAPDH (anti-GAPDH) in the input and IP fractions. GAPDH served as a loading control for the input fractions and a negative control in the IP experiments. **(c)** *Separation of proteins associated with the WT and mutant ORF1p-FLAG proteins.* The WT and M8 (RBM) mutant ORF1p-FLAG IP complexes were separated by SDS-PAGE using a 4-15% gradient gel and silver staining visualized the proteins. Protein size standards (kDa) are shown at the left of the gel. Black arrowhead, the expected molecular weight of ORF1p-FLAG. **(d)** *Gene Ontology (GO) analysis identifies cellular proteins enriched in IP WT ORF1p-FLAG vs. the mutant ORF1p-FLAG complex.* Cellular proteins present in the WT ORF1p and (M8 [RBM])-FLAG mutant IP complexes were identified using LC-MS/MS. Proteins having at least five peptide matches to the UniProt database (https://www.uniprot.org/) were subjected to PANTHER statistical enrichment analysis. The top 10 GO terms with the lowest false discovery rates (FDRs) are sorted in descending values. X-axis, −log10 FDR. Y-axis, GO term. Red lettering, viral related GO terms. **(e)** *Leading Edge Analysis identifies interferon-related gene sets enriched upon WT ORF1p-FLAG immunoprecipitation.* Gene Set Enrichment Analysis (GSEA) of peptides immunoprecipitated in WT ORF1p-FLAG *vs.* (M8 [RBM])-FLAG IP complexes was performed using hallmark gene sets in the Molecular Signatures Database (<u>MSigDB:</u> www.gsea-msigdb.org/gsea/msigdb/), followed by Leading Edge Analysis to determine gene set enrichment scores. The top five hallmark gene sets with the highest normalized enrichment score (NES) are sorted in descending values. X-axis, NES. Y-axis, hallmark gene sets. **(f)** *The expression of engineered L1s modestly up-regulates IFN-α expression.* HEK293T were transfected with either pCEP4 (an empty vector control), pJM101/L1.3FLAG (WT), pJM105/L1.3 (RT-), or pALAF008 (M8 [RBM]). RT-qPCR was used to quantify IFN-α (primer set: IFN-α) and L1 expression (primer set: *mneoI* [Alu or L1]) ~96 hours post-transfection. IFN-α and L1 expression levels were normalized using β-actin (*ACTB*) as a control (primer set: Beta-actin). X-axis, name of constructs. Control, pCEP4. Y-axis, relative RNA expression levels normalized to the pCEP4 empty vector control. Red bars, normalized IFN-α expression levels. Gray bars, normalized L1 expression levels. Values from three independent biological replicates ± SEM are depicted in the graph. The *p*-values were calculated using a one-way ANOVA followed by Bonferroni-Holm post-hoc tests: pairwise comparisons of IFN-α relative to the pCEP4 control, *p* = 0.00028*** (WT); 0.00011*** (RT-); 3.14 × 10^−6^*** (M8 [RBM]). Pairwise comparisons of IFN-α: WT *vs.* RT-, *p* = 0.21^ns^; WT *vs.* M8 (RBM), p = 0.00036***. Pairwise comparisons of L1 relative to WT, *p* = 0.87^ns^ (RT-), *p =* 0.10^ns^ (M8 [RBM]); ns: not significant; *** *p*<0.001.

### Immune-related proteins associate with the WT ORF1p complex

To identify cellular proteins that differentially interact with the WT ORF1p-FLAG and M8/RBM-FLAG protein complexes, we conducted immunoprecipitation followed by liquid chromatography-tandem mass spectrometry (IP/LC-MS/MS) analyses (**Fig. 2c; see Source Data.xlsx file**). Proteins that exhibited five or more peptide matches to the UniProt database (https://www.uniprot.org/) then were subjected to gene ontology (GO) analysis. “Viral transcription” was the most statistically significant enriched PANTHER GO term with the lowest false discovery rate (FDR) identified in the WT ORF1p-FLAG IP/LC-MS/MS experiments (**Fig. 2d and Supplementary Table 1**); “viral gene expression” represented the third most enriched GO term (**Fig. 2d**). Analysis of the M8/RBM-FLAG IP/LC-MS/MS data did not return any significant GO term enrichments. Thus, there was a significant enrichment of proteins encoded by viral process-related genes in the WT ORF1p-FLAG *vs.* the M8/RBM-FLAG IP/LC-MS/MS analyses (**Fig. 2d**; FDR<0.05).

We next performed Gene Set Enrichment Analysis (GSEA) followed by leading edge analysis to determine if there was an enrichment of hallmark gene sets (Molecular Signatures Database [MsigDB]) from the proteins identified in the WT ORF1p-FLAG *vs*. M8/RBM-FLAG IP/LC-MS/MS experiments (**see Methods**). These analyses identified two interferon-related gene sets (interferon gamma and interferon alpha responses), which exhibited normalized enrichment score (NES) of ~1.6 and ~1.4, respectively, among the top five most significantly enriched gene sets in the WT ORF1p-FLAG *vs.* that M8/RBM-FLAG data (**Fig. 2e and Supplementary Table 2, see Methods**).

The overexpression of engineered L1s previously was reported to modestly induce the type I IFN response^42,45,46,62^. Thus, we tested whether there was a difference in IFN-α induction in HEK293T cells transfected with either pJM101/L1.3FLAG (WT ORF1p-FLAG), pJM105/L1.3 (reverse transcriptase deficient [RT-]), or pALAF008_L1.3FLAG_M8 (M8/RBM-FLAG). Expression of the WT ORF1p-FLAG or RT-deficient mutant construct each led to a moderate induction (~2.5-fold increase) of IFN-α transcription (**Fig. 2f**). By comparison, M8/RBM-FLAG expression induced a more significant (~4-fold increase) in IFN-α transcription, when compared to a mock control (**Fig. 2f**). Notably, the L1 RNA levels of the mutants were similar to the WT L1 (using a primer set that amplified the *mneoI* retrotransposition reporter cassette) (**Fig. 2f**). Because the expression of each construct upregulates IFN-α expression, the data suggest that L1 RNA, but not L1 cDNA or L1 retrotransposition *per se*, are responsible for the modest induction of type I IFN expression.

### Proteins produced by Interferon-Stimulated Genes (ISGs) as potential L1 regulators

A number of proteins expressed from interferon stimulated genes (ISGs) have been reported to influence L1 and/or Alu retrotransposition. These proteins include: (1) MOV10, an RNA helicase^53,63,64^; (2) ADAR1, a double-stranded RNA-specific adenosine deaminase^65^; (3) APOBEC3A, 3B, 3C, 3F, and, for Alu, APOBEC3G, paralogs of the apolipoprotein B editing complex enzyme catalytic polypeptide-like 3 containing cytidine deaminase activity^66–73^; (4) TREX1, a three prime repair exonuclease 1^41,74,75^; (5) ZAP, a zinc-finger antiviral protein^50,54^; (6) SAMHD1, a sterile alpha motif (SAM) domain and histidine-aspartate (HD) domain-containing protein 1^76–78^; (7) RNase H2^79,80^; and (8) RNaseL, a protein that is activated by 2’,5’-oligoadenylate (2-5A) synthetase (OAS) to enzymatically degrade L1 RNA^81^. Thus, we hypothesized that the ISG proteins associated with L1 RNPs may directly regulate L1 retrotransposition and/or L1-mediated IFN-α expression.

To test the above hypothesis, we screened the top 300 proteins that associated with WT ORF1p-FLAG in our IP/LC-MS/MS analyses using the interferome database (www.interferome.org)^82^. We used a strict threshold to identify proteins that exhibited a >10-fold increase in expression upon type I IFN induction, leading to the identification of seven proteins. Two proteins, ADAR1 and ZAP, previously were reported to inhibit L1 retrotransposition^50,54,65^. We reasoned the other five proteins, DDX60L, HELZ2, HERC5, IFIT1 and OASL, might also be involved in the regulation of L1 retrotransposition (**Fig. 3a**). Notably, these proteins were enriched in the WT ORF1-FLAG IP/LC-MS/MS data *vs.* M8/RBM-FLAG by at least 2-fold (**Fig. 3b**).

**Figure. 3:**
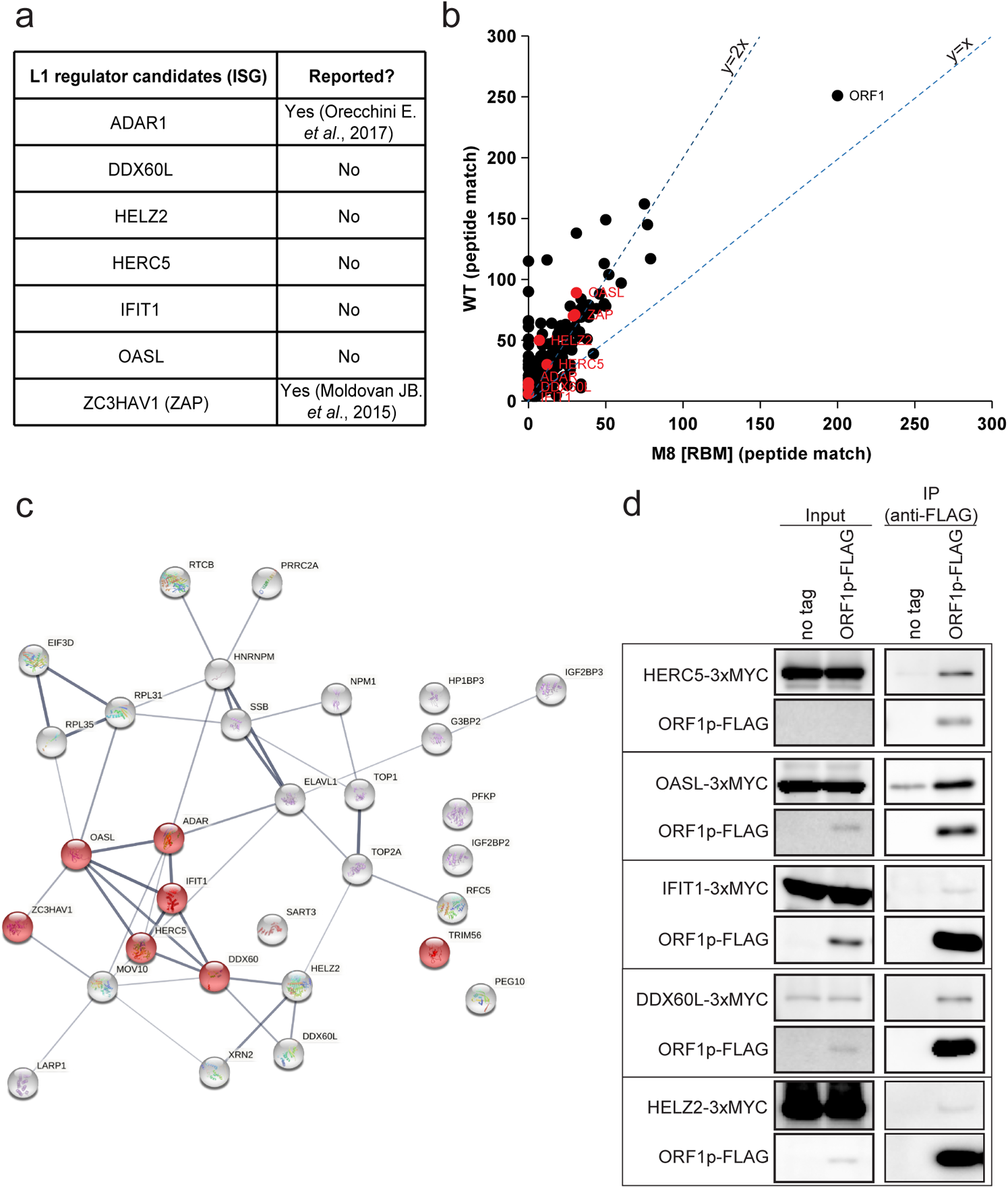
A network of ISGs that potentially affect WT L1 retrotransposition. **(a)** *ISG candidate proteins that may affect L1 biology*. The top 300 proteins identified in the WT ORF1p-FLAG complex were analyzed against the interferome database (http://www.interferome.org/interferome/home.jspx) to identify proteins that exhibit a >10-fold increase in expression upon type I interferon induction. ADAR1 and ZAP previously were reported to inhibit L1 retrotransposition; DDX60L, HELZ2, HERC5, IFIT1, and OASL represent candidate ISG proteins that may play a role in L1 biology. **(b)** *Scatter plot analysis of the top 300 proteins identified in the WT ORF1p-FLAG and ORF1p-FLAG (M8 [RBM]) mutant IP complexes*. X-axis, the number of matching peptides to proteins in the UniProt database found in the ORF1p-FLAG (M8 [RBM]) mutant IP complex. Y-axis, the number of matching peptides to proteins in the UniProt database in the WT ORF1p-FLAG IP complex. Red dots, the proteins enriched in the WT ORF1p-FLAG IP complexes listed in panel (a). **(c)** *String database analysis of WT ORF1-FLAG associated proteins*. Proteins identified in the WT ORF1p-FLAG complex that exhibited a >5-fold increase in expression upon type I IFN induction were subjected to String analysis. Red spheres, proteins annotated as antiviral defense proteins in UniProt. Thickness of the inter-connecting lines, the strength of association based on the number of independent channels supporting the putative interactions. **(d)** *Independent confirmation that ISG proteins interact with WT ORF1p-FLAG.* HEK293T cells were co-transfected with either pJM101/L1.3 (no tag) or pJM101/L1.3-FLAG (ORF1p-FLAG) and the following individual carboxyl-terminal 3xMYC epitope-tagged ISG expression vectors: pALAF015 (HELZ2), pALAF016 (IFIT1), pALAF021 (DDX60L), pALAF022 (OASL), or pALAF023 (HERC5). The input and anti-FLAG IP reactions were analyzed by western blotting using an anti-FLAG (to detect ORF1p-FLAG) or an anti-MYC (to detect ISG proteins) antibody.

We next relaxed our threshold and screened the interferome database for WT ORF1p-FLAG associated proteins exhibiting a 5-fold increase in expression upon type I IFN induction (**Supplementary Table 3**) and then used StringDB (https://string-db.org/)^83^ to test for possible associations among the putative type I IFN interferon inducible proteins that preferentially associated with WT ORF1p-FLAG^45,46^. Most of the ISG candidates exhibiting a >10-fold increase in expression upon type I IFN induction (*i.e.*, ADAR1, ZAP, DDX60L, HELZ2, HERC5, IFIT1 and OASL), with the exception of HELZ2 and DDX60L, were annotated as antiviral defense proteins in UniProt (https://www.uniprot.org/) (**Fig. 3c**, red circles; FDR, 4.5×10^−8^, interaction strength, 1.55). Thus, a network of antiviral ISG proteins may regulate L1 RNA and/or L1 RNP dynamics.

To validate the interaction of proteins identified in the above analyses with WT ORF1p-FLAG, we conducted additional co-IP experiments. Briefly, pJM101/L1.3FLAG (WT ORF1p-FLAG) was co-transfected into HEK293T cells with individual ISG protein expression vectors (HELZ2, IFIT1, DDX60L, OASL, and HERC5) containing three copies of a MYC epitope tag at their respective carboxyl-termini. An anti-FLAG primary antibody then was used to immunoprecipitate associated proteins from HEK293T whole cell extracts and an anti-MYC antibody was used to confirm associations between WT ORF1p-FLAG and the candidate ISG proteins. WT ORF1p-FLAG co-immunoprecipitated HERC5, OASL, IFIT1, DDX60L, and HELZ2 (**Fig. 3d**).

### The ISG proteins, HELZ2, OASL, and HERC5 inhibit L1 retrotransposition

To determine whether ectopic overexpression of the identified ISG proteins affect L1 retrotransposition, we co-transfected HeLa-JVM or HEK293T cells with a WT human L1 expression construct containing either a *mblastI* (pJJ101/L1.3) or *mEGFPI* (cepB-gfp-L1.3) retrotransposition indicator cassette and the carboxy-terminal MYC epitope-tagged HELZ2, IFIT1, DDX60L, OASL, or HERC5 expression vectors. L1 retrotransposition efficiencies then were determined by counting the resultant number of blasticidin-resistant foci or EGFP-positive cells (**Fig. 1c, see Methods**). A MOV10 expression vector, also containing a carboxyl-terminal 3x MYC epitope tag served as a positive control. The overexpression of DDX60L and IFIT1 did not significantly inhibit L1 retrotransposition in HeLa-JVM (**Figs. 4a and 4b**) or HEK293T cells (**Supplementary Figs. 4a and 4b**), although we note the expression of DDX60L was barely detected by western blot in either cell line (**Fig. 4b and Supplementary Fig. 4b**). By comparison, overexpression of HERC5, HELZ2, and OASL reduced retrotransposition by at least 2-fold in the *mblastI*-based L1 retrotransposition assay conducted in HeLa-JVM cells (**Fig. 4a**) and by ~90% in the *mEGFPI*-based L1 retrotransposition assay conducted in HEK293T cells (**Supplementary Fig. 4a**).

**Figure. 4:**
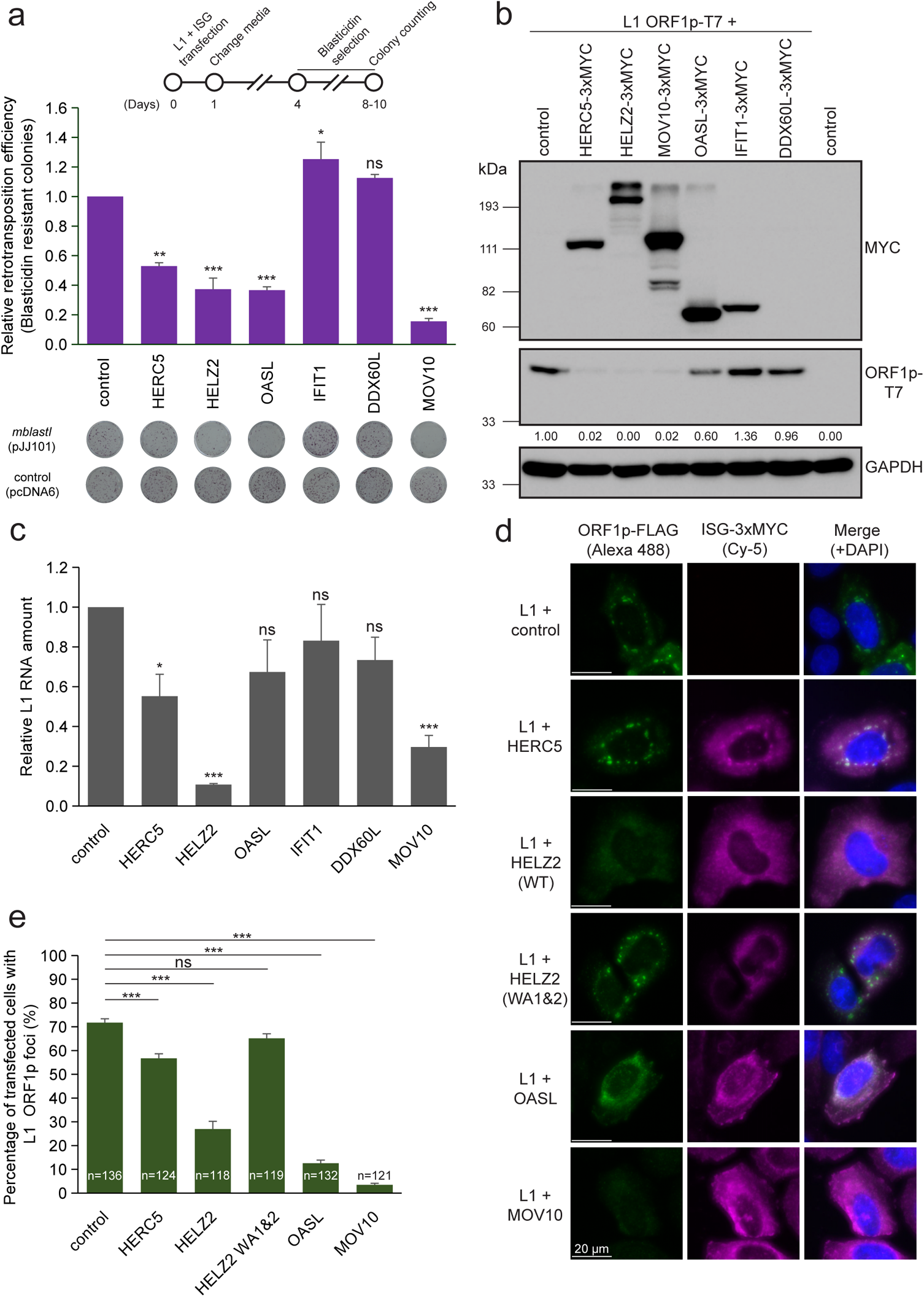
A subset of ISG proteins affect steady state L1 RNA levels, ORF1p cytoplasmic foci formation, and/or L1 retrotransposition. **(a)** *Overexpression of HERC5, HELZ2, or OASL inhibit L1 retrotransposition.* HeLa-JVM cells were co-transfected with pJJ101/L1.3, which contains the *mblastI* retrotransposition indicator cassette, and either pCMV-3Tag-8-Barr or one of the following carboxyl-terminal 3xMYC epitope-tagged ISG protein expression plasmids: pALAF015 (HELZ2), pALAF016 (IFIT1), pALAF021 (DDX60L), pALAF022 (OASL), pALAF023 (HERC5), or pALAF024 (MOV10) according to the timeline shown at the top of the figure. A blasticidin expression vector (pcDNA6) was co-transfected into cells with either pCMV-3Tag-8-Barr or an individual ISG protein expression plasmid (see plates labeled control [pcDNA6]) to assess cell viability. The retrotransposition efficiencies then were normalized to the respective toxicity control. X-axis, name of the control (pCMV-3Tag-8-Barr) or ISG protein expression plasmid. Y-axis, relative retrotransposition efficiency normalized to the pJJ101/L1.3/pCMV-3Tag-8-Barr co-transfected control. Representative results of the retrotransposition (see plates labeled *mblastI* [pJJ101]) and toxicity (see plates labeled *control* [pcDNA6]) assays are shown below the graph. Pairwise comparisons relative to the pJJ101/L1.3 + pCMV-3Tag-8-Barr control: *p* = 8.0 × 10^−5^** (HERC5); 4.4 × 10^−6^*** (HELZ2); 4.9 × 10^−6^*** (OASL); 0.011* (IFIT1); 0.12^ns^ (DDX60L); and 1.7 × 10^−7^*** (MOV10). MOV10 served as a positive control in the assay. **(b)** *Expression of the ISG proteins in HeLa-JVM cells.* HeLa-JVM cells were co-transfected with pTMF3, which expresses a version of ORF1p containing a T7 gene 10 carboxyl epitope tag (ORF1p-T7), and either a pCMV-3Tag-8-Barr (control) or the individual ISG-expressing plasmids used in panel (a). Whole cell extracts were subjected to western blot analysis 48 hours post-transfection. ISG proteins were detected using an anti-MYC antibody. ORF1p was detected using an anti-T7 antibody. GAPDH served as a loading control. The relative band intensities of ORF1p-T7 are indicated under the ORF1p-T7 blot; they were calculated using ImageJ software and normalized to the respective GAPDH bands. **(c)** *HELZ2 expression leads to a reduction in the steady state level of L1 RNA.* HeLa-JVM cells were transfected as in panel (b). L1 RNA levels were determined by performing RT-qPCR using a primer set specific to RNAs derived from the transfected L1 (primer set: L1 [SV40]) and then were normalized to *ACTB* RNA levels (primer set: Beta-actin). X-axis, name of the constructs. Y-axis, relative level of L1 RNA normalized to the ORF1-T7 + pCMV-3Tag-8-Barr control. Pairwise comparisons relative to the control: *p* = 0.032* (HERC5); 1.7 × 10^−5^*** (HELZ2): 0.14^ns^ (OASL); 0.29^ns^ (IFIT1); 0.20^ns^ (DDX60L); and 4.4 × 10^−4***^ (MOV10). **(d)** *Differential effects of ISG proteins on ORF1p-FLAG cytoplasmic foci formation*. HeLa-JVM cells were co-transfected with pJM101/L1.3FLAG (WT ORF1p-FLAG) and either a pCEP4 empty vector (control) or one of the following carboxyl-terminal 3xMYC epitope-tagged ISG protein expression plasmids: pALAF015 (HELZ2); pALAF027 (HELZ2 WA1&2); pALAF022 (OASL); pALAF023 (HERC5); or pALAF024 (MOV10) to visualize WT ORF1p-FLAG cytoplasmic foci and co-localization between WT ORF1p-FLAG and the candidate ISG protein. Shown are representative fluorescent microscopy images. White scale bars, 20 µm. **(e)** *Quantification of L1 cytoplasmic foci formation.* X-axis, name of the constructs co-transfected with pJM101/L1.3FLAG (WT ORF1p-FLAG); control, pCEP4. Y-axis, percentage of transfected cells with visible ORF1p signal exhibiting ORF1p-FLAG cytoplasmic foci. The numbers (n) within the green rectangles indicate the number of analyzed cells in each experiment. Pairwise comparisons relative to the pJM101/L1.3FLAG (WT ORF1p-FLAG) + pCEP4 control: *p* = 8.6 × 10^−4^*** (HERC5); 1.2 × 10^−7^*** (HELZ2); 0.098^ns^ (HELZ2 WA1&2); 1.0 × 10^−10^*** (OASL); 2.7 × 10^−9^*** (MOV10). Values represent the mean ± SEM from three (in panels [a] and [e]) or six (in panel [c]) independent biological replicates. The *p*-values were calculated using one-way ANOVA followed by Bonferroni-Holm post-hoc tests; ns: not significant; * *p<*0.05; ** *p*<0.01; *** *p<*0.001.

### Some ISG proteins affect ORF1p and L1 mRNA levels

To further understand how ISG proteins might inhibit L1 retrotransposition, we co-transfected a full-length RC-L1 (pTMF3) and either the HELZ2, IFIT1, DDX60L, OASL, HERC5, or MOV10 expression vectors into HeLa-JVM or HEK293T cells and examined whether the ISG proteins affected ORF1p and/or L1 RNA expression. Western blot analysis revealed a similar data trend in HeLa-JVM and HEK293T cells: the steady state ORF1p levels were significantly decreased by co-expression of HERC5, HELZ2, and MOV10, were modestly reduced by the co-expression of OASL, but were not changed by the co-expression of IFIT1 or DDX60L (**Fig. 4b and Supplementary Fig. 4b**). RT-qPCR analyses, using a probe set that specifically recognizes the SV40 polyA signal of the plasmid-expressed L1 RNA, revealed that HELZ2 significantly reduced L1 RNA levels in HeLa-JVM cells (**Fig. 4c**, ~90% reduction of the WT L1 control). MOV10 co-expression resulted in a ~70% reduction in L1 RNA when compared to the WT L1 control, which is consistent with previous reports^64,84^.

We next tested whether the co-transfection of pJM101/L1.3FLAG (WT ORF1p-FLAG) with the individual ISG protein expression vectors (*i.e.*, HELZ2, HERC5, OASL, and MOV10) affected ORF1p-FLAG cytoplasmic foci formation in HeLa-JVM cells (**Figs. 4d and 4e**). Greater than 70% of transfected cells expressing WT ORF1p-FLAG exhibited cytoplasmic foci (**Fig. 4e**), which is consistent with previous results^49^ (**see Supplementary Fig. 3d**). Co-expression of HERC5 did not dramatically affect ORF1p cytoplasmic foci formation in HeLa-JVM cells (**Fig. 4e**, ~55% of cells contained ORF1p cytoplasmic foci that associated with HERC5). By comparison, the co-expression of HELZ2, OASL, and MOV10 resulted in a decrease in ORF1p-FLAG cytoplasmic foci (**Fig. 4e**, ~30%, ~15%, and ~5% of cells, respectively) and very few of these foci associated with the relevant ISG protein (**Fig. 4e**). In aggregate, these data suggest: (1) HERC5 destabilizes ORF1p, but does not affect its cellular localization; (2) OASL mainly impairs ORF1p cytoplasmic foci formation; and (3) HELZ2 reduces the levels of L1 RNA, ORF1p, and ORF1p cytoplasmic foci formation. Thus, different ISGs appear to affect different steps of the L1 retrotransposition cycle.

### The HELZ2 helicase activity is important for inhibition of L1 retrotransposition

HELZ2 contains two putative helicase domains (helicase 1 and helicase 2) that flank a putative exoribonuclease RNase II/R (RNB) domain (**Fig. 5a and Supplementary Fig. 5a**). Because proteins containing a RNB domain often possess 3’ to 5’ single-strand exoribonuclease activity^85,86^, we aligned the protein sequences of RNB-containing proteins from human, yeast and *E. coli* to identify evolutionarily conserved aspartic acid residues, which when mutated, are predicted to impair exoribonuclease activity^85–87^ (**Supplementary Fig. 5b**). We mutated three conserved aspartic acid residues in HELZ2 to asparagine residues (D1346N/D1354N/D1355N) and assayed whether this triple mutant affects L1 retrotransposition. This triple mutant generally only had minor effects (*i.e.*, less than 2-fold) on L1 retrotransposition efficiency in HeLa-JVM and HEK293T cells when compared to the WT HELZ2 control (**Supplementary Figs. 5c and 5d**).

**Figure. 5:**
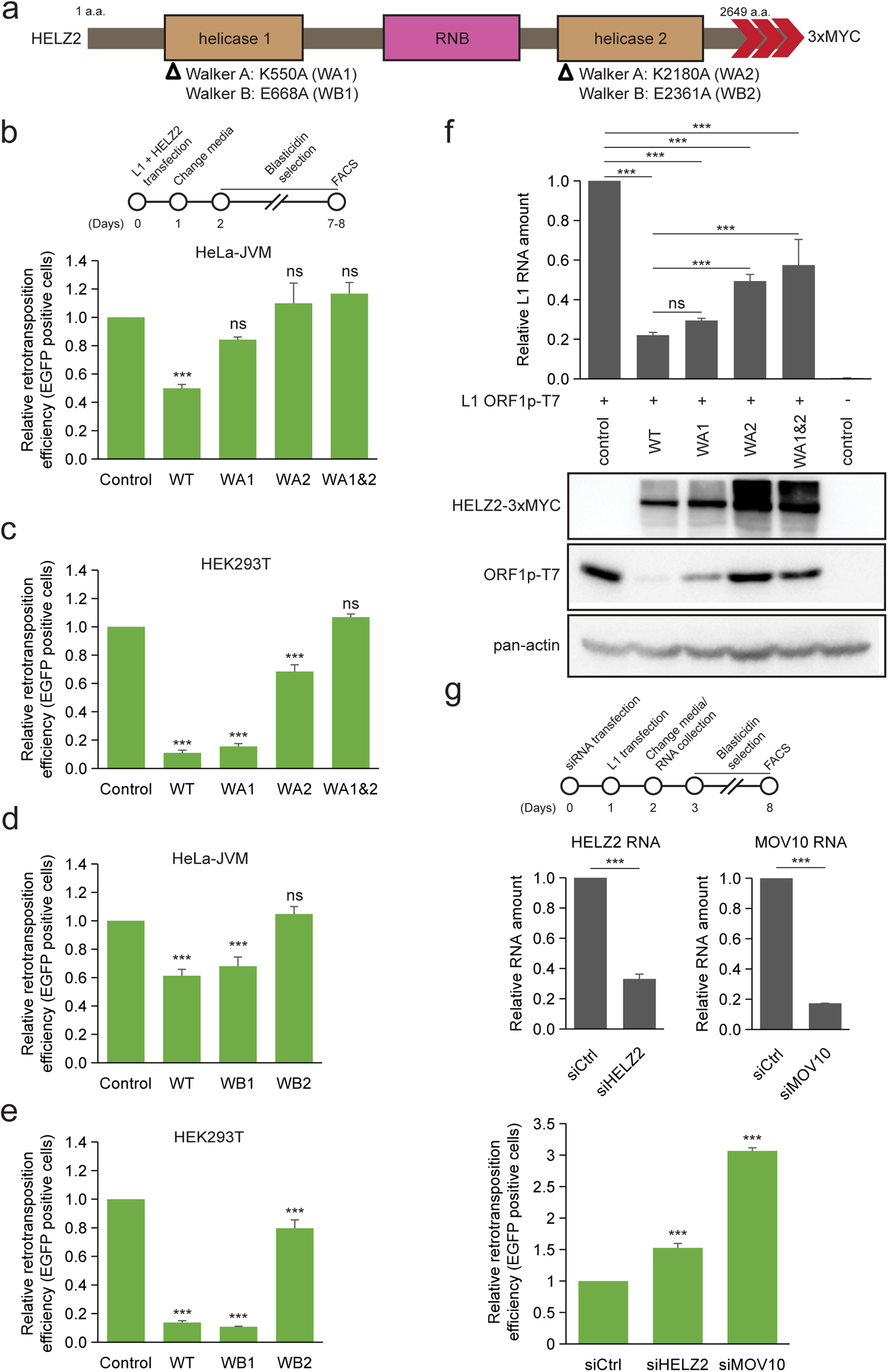
The HELZ2 helicase activity is critical for L1 inhibition. **(a)** *Schematic of the HELZ2 protein domains.* HELZ2 contains two putative helicase domains (helicase 1 and helicase 2), which surround a putative RNB exonuclease domain. Open triangles, positions of missense mutation in conserved amino acids within the Walker A (WA) and Walker B (WB) boxes in the helicase 1 and helicase 2 domains: K550A (WA1); K2180A (WA2); E668A (WB1); and K2361A (WB2). Red arrowheads, relative positions of the 3xMYC carboxyl-terminal epitope tag in the HELZ2 expression constructs. **(b)** *The effect of mutations in the Walker A box on L1 retrotransposition.* HeLa-JVM cells were co-transfected with cepB-gfp-L1.3, which contains a *mEGFPI* retrotransposition indicator cassette, and either pCMV-3Tag-8-Barr (control), pALAF015 (WT HELZ2), or one of the following HELZ2 expression plasmids that contain a mutation(s) in the Walker A box: pALAF025 (WA1); pALAF026 (WA2); or pALAF027 (WA1&2). Cells co-transfected with cepB-gfp-L1.3RT(−) intronless and either pCMV-3Tag-8-Barr, pALAF015 (WT HELZ2), or a mutant HELZ2 plasmid served as transfection normalization and toxicity controls. Top, timeline of the assay for panels (b), (c), (d), and (e). X-axis, name of HELZ2 expression constructs co-transfected into cells with cepB-gfp-L1.3; control, pCMV-3Tag-8-Barr. Y-axis, relative retrotransposition efficiency normalized to the cepB-gfp-L1.3 + pCMV-3Tag-8-Barr control. Pairwise comparisons relative to the control: *p* = 0.00087*** (WT HELZ2); 0.26^ns^ (WA1); 0.32^ns^ (WA2); and 0.32^ns^ (WA1&2). **(c)** *The effect of mutations in the Walker A box on L1 retrotransposition in HEK293T cells*. HEK293T cells were co-transfected as in panel (b). Retrotransposition efficiencies were calculated as described in panel (b). Pairwise comparisons relative to the cepB-gfp-L1.3 (*mEGFPI*) + pCMV-3Tag-8-Barr control: *p* = 2.5 × 10^−11^*** (WT HELZ2); 3.5 × 10^−11^*** (WA1); 1.7 × 10^−6^*** (WA2); and 0.070^ns^ (WA1&2). **(d)** *The effect of mutations in the Walker B box on L1 retrotransposition in HeLa-JVM cells*. L1 retrotransposition assays were performed as in panel (b). Co-transfections were performed using an individual HELZ2 expression plasmid containing a mutation in the Walker B box: pALAF028 (WB1) or pALAF029 (WB2). Retrotransposition efficiencies were calculated as described in panel (b). Pairwise comparisons relative to the L1.3 + pCMV-3Tag-8-Barr control: *p* = 9.5 × 10^−5^*** (WT); 0.0004*** (WB1); and 0.43^ns^ (WB2). **(e)** *The effect of mutations in the Walker B box on L1 retrotransposition in HEK293T cells*. HEK293T cells were co-transfected as in panel (b). Retrotransposition efficiencies were calculated as described in panel (b). Pairwise comparisons relative to the cepB-gfp-L1.3 (mEGFPI) + pCMV-3Tag-8-Barr control: *p* = 9.4 × 10^−10^*** (WT); 8.4 × 10^−10^*** (WB1); and 8.7 × 10^−4^*** (WB2). **(f)** *Mutations in the HELZ2 helicase domains reduce the ability to inhibit L1 ORF1p and RNA.* HeLa-JVM cells were transfected with pTMF3 (L1 ORF1p-T7), denoted by + symbol, and either pCMV-3Tag-8-Barr (control), pALAF015 (WT HELZ2), or an individual HELZ2 expression plasmid containing a mutation(s) in the Walker A box: pALAF025 (WA1), pALAF026 (WA2), or pALAF027 (WA1&2). Top: L1 RNA levels were determined by RT-qPCR using primers directed against sequences in the transfected L1 RNA (primer set: L1 [SV40]) and then were normalized to *ACTB* RNA levels (primer set: Beta-actin). Pairwise comparisons relative to the pTMF3 (L1 ORF1p-T7) + pCMV-3Tag-8-Barr control: *p* = 9.5 × 10^−9^*** (WT); 1.9 × 10^−8^*** (WA1); 7.3 × 10^−7^*** (WA2); and 1.5 × 10^−6^*** (WA1&2). Pairwise comparisons relative to the pTMF3 (L1 ORF1p-T7) + WT HELZ2: *p* = 0.56^ns^ (WA1); 5.9 × 10^−4^*** (WA2); 1.9 × 10^−4^*** (WA1&2). Bottom: western blot image displaying ORF1p-T7 bands. HELZ2 expression was detected using an anti-MYC antibody. ORF1p was detected using an anti-T7 antibody. Pan-actin served as a loading control. **(g)** *Short-interfering RNA (siRNA)-mediated knockdown of endogenous HELZ2 increases L1 retrotransposition.* Top, timeline of the assay conducted in HeLa-JVM cells. Cells were transfected with a non-targeting siRNA control (siCtrl), siRNA targeting HELZ2 (siHELZ2), or siRNA targeting MOV10 (siMOV10). Middle left panel, HELZ2 RNA levels in siRNA treated cells. Middle right panel, MOV10 RNA levels in siRNA treated cells. X-axes, name of the siRNA. HELZ2 and MOV10 RNA levels were determined using RT-qPCR (primer sets: HELZ2 and MOV10, respectively) and then were normalized to *ACTB* RNA levels (primer set: Beta-actin). Y-axes, relative HELZ2 or MOV10 RNA levels normalized to the siCtrl. A two-tailed, unpaired Student’s t-test was used to calculate the *p*-values relative to the siRNA control: *p* = 3.1 × 10^−5^*** (siHELZ2); and 5.2 × 10^−5^*** (siMOV10). Bottom panel, HeLa-JVM cells were transfected with either siCtrl, siHELZ2, or siMOV10, followed by transfection with either cepB-gfp-L1.3 or cepB-gfp-L1.3RT(−) intronless, which was used to normalize transfection efficiencies. X-axis, name of the siRNA. Y-axis, relative retrotransposition efficiency. Pairwise comparisons relative to the non-targeting siRNA control: *p* = 2.9 × 10^−4^*** (siHELZ2); and 2.0 × 10^−7^*** (siMOV10). All the reported values represent the mean ± SEM from three independent biological replicates. The *p*-values, except for the RT-qPCR experiment shown in panel (g), were calculated using a one-way ANOVA followed by a Bonferroni-Holm post-hoc tests. ns: not significant; * *p<*0.05; *** *p<*0.001.

We next tested whether mutations in the putative HELZ2 helicase domains affect L1 retrotransposition. We mutated conserved amino acids in the Walker A and Walker B boxes thought to be required for ATP binding (WA1 [K550A] and WA2 [K2180A]) or ATP hydrolysis (WB1 [E668Ap] and WB2 [E2361A]), respectively^88–90^. The WA1 mutant demonstrated a low, but not statistically significant decrease in L1 retrotransposition efficiency in HeLa-JVM cells (**Fig. 5b**), but exhibited a significant decrease in L1 retrotransposition in HEK293T cells (**Fig. 5c**). The WA2 did not significantly inhibit L1 retrotransposition in HeLa-JVM cells (**Fig. 5b**), but showed a low level of inhibition in HEK293T cells (**Fig. 5c**). The WA1&2 double mutant was unable to inhibit L1 retrotransposition in either HeLa-JVM or HEK293T cells (**Figs. 5b and 5c**). By comparison, the WB1 mutant retained the ability to inhibit L1 retrotransposition in HeLa-JVM and HEK293T cells (**Figs. 5d and 5e, respectively**), whereas the WB2 mutant did not inhibit L1 retrotransposition in HeLa-JVM cells (**Fig. 5d**) and only exhibited minor inhibition in HEK293T cells (**Fig. 5e**). In general, the WA2 and WB2 mutants consistently exhibited a less severe inhibition of L1 retrotransposition when compared to WA1 and WB1 mutants, suggesting the importance of the helicase 2 domain in the inhibition of retrotransposition.

Additional experiments revealed that the WA1 mutant reduced both ORF1p-T7 and L1 RNA levels in HeLa-JVM cells (**Fig. 5f**); the WA2 and WA1&2 double mutant partially reduced L1 RNA levels in comparison to the WT control, but did not affect the steady state levels of the ORF1p-T7 protein (**Fig. 5f**). Importantly, we did not observe a noticeable reduction in the steady state levels of the HELZ2 mutant proteins (**Fig. 5f**, bottom panel), suggesting that the effects on L1 retrotransposition are not due to mutant HELZ2 protein instability (**Fig. 5f**). Finally, the WA1&2 double mutant did not affect the ability of ORF1p-FLAG to localize to cytoplasmic foci when compared to WT HELZ2 (**Figs. 4d and 4e**). A union of the above data suggest that the HELZ2 helicase activity has a more pronounced effect than the HELZ2 RNase activity on L1 retrotransposition and that mutations in the HELZ2 helicase domains affect L1 RNA stability, ORF1p levels, and ORF1p cytoplasmic localization to different extents.

### Knockdown of endogenous HELZ2 enhances L1 retrotransposition

To determine whether endogenous HELZ2 could inhibit L1 retrotransposition, we used small interfering RNAs (siRNAs) to reduce HELZ2 and MOV10 levels in HeLa-JVM cells. Control RT-qPCR experiments revealed a ~70% and ~80% knockdown of HELZ2 and MOV10 RNAs, respectively, when compared to a non-targeting siRNA control (**Fig. 5g**, middle panel); *mEGFPI*-based assays revealed a ~1.5-fold and ~3-fold increase in L1 retrotransposition efficiency in the siHELZ2 and siMOV10 treated cells, respectively (**Fig. 5g**, bottom panel). Thus, endogenous HELZ2 may also suppress L1 retrotransposition.

### HELZ2 recognizes L1 RNA independent of RNP formation

We further investigated the mechanism of association between ORF1p-FLAG and HELZ2. Treatment of the ORF1p RNP complex with RNase A abolished the ORF1p-FLAG/HELZ2 interaction, suggesting that HELZ2, like PABPC1, associates with ORF1p in an RNA-dependent manner^50,61^ (**Fig. 6a**).

**Figure. 6:**
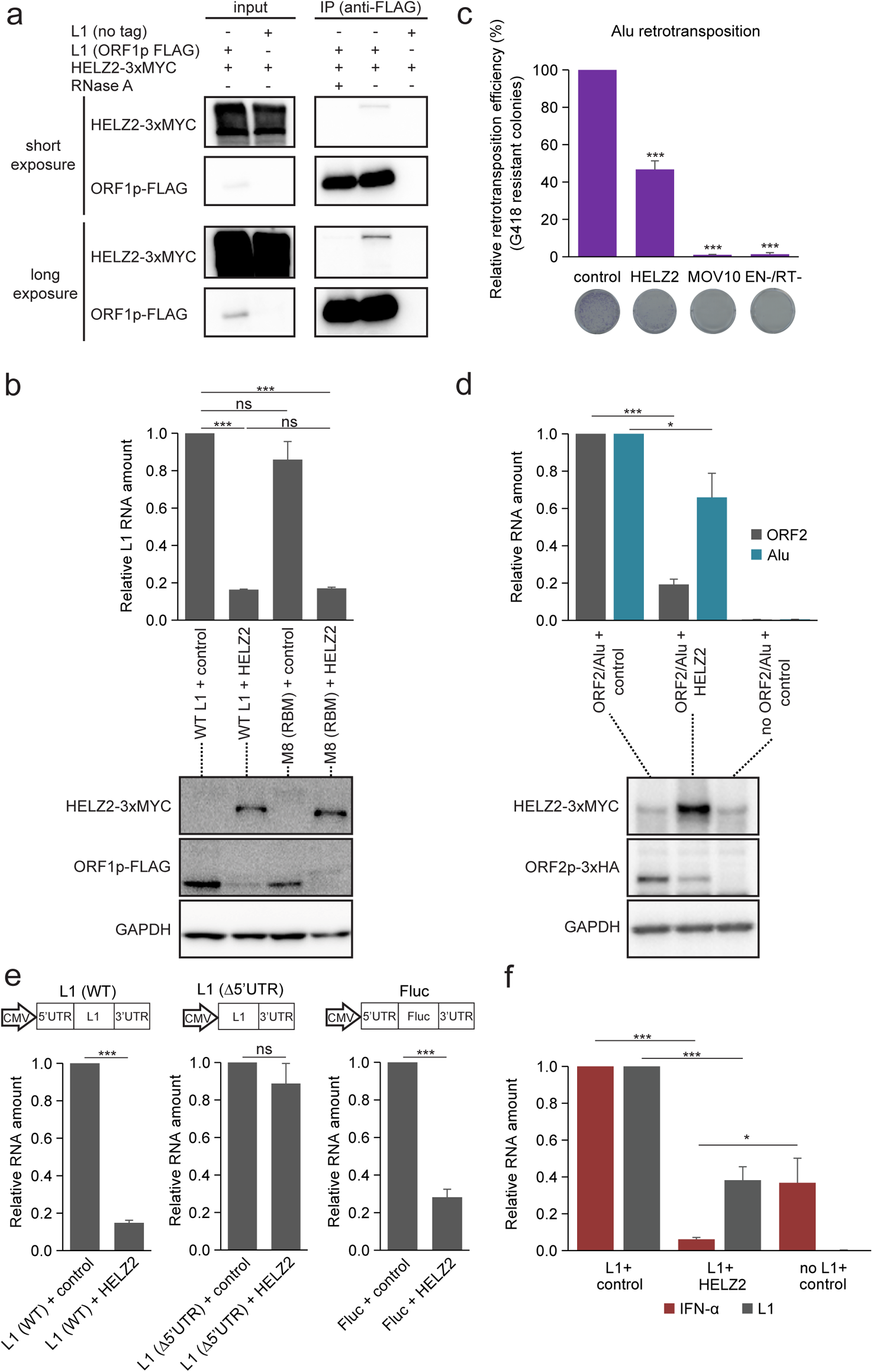
HELZ2 destabilizes L1 RNA through recognition of the L1 5’UTR sequence, leading to attenuation of L1-mediated IFN-α induction. **(a)** *The association between ORF1p and HELZ2 is RNA-dependent*. HEK293T cells were co-transfected with pALAF015 (HELZ2-3xMYC) and either pJM101/L1.3FLAG (WT ORF1p-FLAG) or pJM101/L1.3 (no tag). The input and anti-FLAG IP fractions were analyzed by western blot using an anti-FLAG antibody to detect ORF1p-FLAG or an anti-MYC antibody to detect HELZ2-3xMYC. Shown are short (top blots) and longer (bottom blots) chemiluminescence western blot exposures. **(b)** *HELZ2 expression reduces steady state levels of L1 RNA and ORF1p independent of ORF1p RNA-binding*. HeLa-JVM cells were co-transfected with pJM101/L1.3FLAG (WT ORF1p-FLAG) or the pALAF008 ORF1p-FLAG (M8 [RBM]) mutant expression plasmid and either pCMV-3Tag-8-Barr (control) or pALAF015 (HELZ2). Top: L1 RNA amounts were determined by RT-qPCR (primer set: L1 [SV40]) and then were normalized to *ACTB* RNA levels (primer set: Beta-actin). The L1 RNA values were normalized to the WT L1 or ORF1p-FLAG (M8 [RBM]) + pCMV-3Tag-8-Barr control transfections. Pairwise comparisons (in parentheses) relative to the (WT L1 + control) are shown: *p* = 7.1×10^−7^*** (WT L1 + HELZ2); 0.090^ns^ (M8 [RBM] + control); 6.7×10^−7^*** (M8 [RBM] + HELZ2). Pairwise comparisons of (WT L1 + HELZ2) *vs*. (M8 [RBM] + HELZ2), *p* = 0.92^ns^. Bottom: ORF1p-FLAG and HELZ2 protein levels were detected by western blot using anti-MYC and anti-FLAG antibodies, respectively. GAPDH served as a loading control. **(c)** *HELZ2 expression inhibits Alu retrotransposition*. HeLa-HA cells were co-transfected with pTMO2F3_Alu (which expresses an Alu element marked with *neo*-based retrotransposition indicator cassette and monocistronic version of L1 ORF2p [see Methods]), pTMO2F3D145AD702A_Alu (which expresses an Alu element marked with *neo*-based retrotransposition indicator cassette and an EN-/RT-mutant version of L1 ORF2 [see Methods]), or phrGFP-C (a transfection normalization control) and either pCMV-3Tag-8-Barr (control), pALAF015 (WT HELZ2), or pALAF024 (WT MOV10). X-axis, name of constructs. Y-axis, the percentage of G418-resistant foci, indicative of Alu retrotransposition, relative to the pTMO2F3_Alu + pCMV-3Tag-8-Barr control (see Methods for more detail). Representative images of G418-resistant foci are shown below graph. Pairwise comparisons relative to the pTMO2F3_Alu + pCMV-3Tag-8-Barr control: *p* = 7.8 × 10^−5^*** (HELZ2); 1.8 × 10^−7^*** (MOV10); and 1.6 × 10^−7^*** (EN-/RT-). **(d)** *HELZ2 expression leads to a reduction in monocistronic ORF2 L1 RNA and ORF2p levels.* HeLa-HA cells were co-transfected with pTMO2H3_Alu (ORF2p-3xHA and Alu) and either pCMV-3Tag-8-Barr (control) or pALAF015 (HELZ2). Top: ORF2 (gray bars) and Alu RNA (blue bars) levels were determined using RT-qPCR (primer sets: L1 [SV40] and mneoI [Alu or L1], respectively) and normalized to *ACTB* RNA levels (primer set: Beta-actin). X-axis, co-transfected constructs name. Y-axis, relative RNA level normalized to the pTMO2H3_Alu (ORF2p-3xHA and Alu) + pCMV-3Tag-8-Barr control. L1 ORF2 RNA pairwise comparison (ORF2/Alu + control *vs.* ORF2/Alu + HELZ2), *p* = 7.2 × 10^−8^***. Alu RNA pairwise comparison (ORF2/Alu + control *vs.* ORF2/Alu + HELZ2), *p* = 0.018*. Bottom: Western blotting using an anti-HA antibody was used to detect ORF2p. GAPDH served as a loading control. **(e)** *The L1 5’UTR is required for HELZ2-mediated reduction of L1 RNA levels.* HeLa-JVM cells were co-transfected with L1 (WT), L1 (Δ5’UTR), or Fluc (a firefly luciferase gene flanked by the L1 5’ and 3’UTRs) and either pCMV-3Tag-8-barr (control) or pALAF015 (HELZ2). Schematics of the constructs are above the bar charts. RNA levels were determined by RT-qPCR using the following primer sets: L1 (SV40) (for L1 WT and L1[Δ5’UTR]) or Luciferase (for Fluc) and then were normalized to *GAPDH* RNA levels (primer set: GAPDH). X-axis, name of respective constructs co-transfected with pCMV-3Tag-8-Barr (control) or pALAF015 (HELZ2); Y-axis, the relative amount of L1 or Fluc-based RNA relative to the relevant pairwise control (*e.g.*, the L1 expression plasmid + pCMV-3Tag-8-Barr or the Fluc-based plasmid + pCMV-3Tag-8-Barr). Two-tailed, unpaired Student’s t-tests: *p* = 3.9 × 10^−7^*** (left plot); 0.35^ns^ (middle plot); 7.1 × 10^−5^*** (right plot). **(f)** *HELZ2 expression represses L1-induced IFN-α expression*. HEK293T cells were co-transfected with pJM101/L1.3FLAG and a pCEP4 empty vector (left: L1 + control); pJM101/L1.3FLAG and pALAF015 (middle: L1 + HELZ2); or only pCEP4 empty vector (right: no L1 + control). IFN-α (red bars) and L1 (gray bars) RNA levels were determined by RT-qPCR (using primer sets [IFN-α] and L1 [FLAG], respectively) and normalized to *ACTB* RNA levels (primer set: Beta-actin). The RNA levels then were normalized to the L1 + pCEP4 transfection (L1 + control). L1 RNA pairwise comparison: (L1 + control *vs.* L1 + HELZ2), *p* = 9.6 × 10^−5^***. IFN-α RNA pairwise comparisons: (L1 + control *vs.* L1 + HELZ2), *p* = 4.0 × 10^−4^***; (L1 + HELZ2 *vs.* no L1 + control), *p* = 0.031*. With the exception of panel (e), values are reported as the mean ± SEM of three independent biological replicates. The *p*-values were calculated using a one-way ANOVA followed by Bonferroni-Holm post-hoc tests. ns: not significant; * *p<*0.05; *** *p<*0.001.

To test whether L1 RNP formation is required for the association between ORF1p-FLAG and HELZ2, we compared the effects of HELZ2 overexpression on L1 RNA and ORF1p protein abundance in HeLa-JVM cells transfected with either pJM101/L1.3FLAG (ORF1p-FLAG) or pALAF008_L1.3FLAG_M8 (M8/RBM-FLAG). RT-qPCR using a probe set that specifically recognizes the SV40 polyA signal of plasmid expressed L1 RNA and western blot experiments conducted with an anti-FLAG antibody revealed a marked reduction in L1 RNA (~80% reduction) and ORF1p levels in both the WT ORF1p-FLAG or M8/RBM ORF1p-FLAG transfected cells upon HELZ2 overexpression when compared to controls (**Fig. 6b**). Thus, HELZ2 overexpression appears to destabilize L1 RNA independent of WT L1 RNP formation.

### HELZ2 overexpression modestly inhibits Alu retrotransposition

To examine whether HELZ2 overexpression affects Alu retrotransposition, HeLa-HA cells^69^ were transfected with an expression plasmid that contains both an engineered Alu-element harboring a *neo*-based retrotransposition indicator cassette (*neo*^Tet^)^91^ and a monocistronic L1 ORF2p-3xFLAG expression cassette^92^. HELZ2 overexpression reduced Alu retrotransposition by ~2-fold when compared to the respective controls (**Fig. 6c**). Additional experiments revealed that HELZ2 overexpression reduced L1 ORF2p and Alu RNA levels by ~80% and ~35%, respectively (**Fig. 6d**); the reduction in L1 RNA levels led to a corresponding decrease L1 ORF2p protein levels (see below). Notably, the reductions in the levels of monocistronic and full-length L1 RNAs upon HELZ2 overexpression were quite similar (*i.e.*, **Fig. 6b *vs.* Fig. 6d**), suggesting that the observed decrease in Alu retrotransposition mainly results from the HELZ2-dependent destabilization of L1 RNA.

### HELZ2 recognizes the 5’UTR of L1 RNA to reduce both L1 RNA levels and IFN-α induction

HELZ2 overexpression adversely affects L1 and Alu retrotransposition. Intriguingly, the sequences of the L1 5’ and 3’UTRs are shared between the full-length L1 and monocistronic ORF2p expression constructs used in these assays. However, the monocistronic ORF2p expression cassette that drives Alu retrotransposition contains a deletion of a conserved polypurine tract (Δppt) in the L1 3’UTR, which does not dramatically affect L1 retrotransposition^17^. Thus, we hypothesized that HELZ2 may recognize either RNA sequences or RNA structures in the L1 5’UTR and/or 3’UTRΔppt to destabilize L1 RNA.

To test the above hypothesis, we deleted the L1 5’UTR sequence from a WT L1 expression construct (pTMF3) that also contains the 3’UTRΔppt sequence and drove L1 expression solely from the cytomegalovirus immediate-early (CMV) promoter (**Fig. 6e**, L1 [Δ5’UTR]; a.k.a. pTMF3_Δ5UTR). As an additional control, we also replaced the L1.3-coding sequences (*ORF1* and *ORF2*) with a firefly luciferase gene in the same WT L1 construct (**Fig. 6e**, Fluc; a.k.a pL1[5&3UTRs]_Fluc). Co-transfection of HeLa-JVM cells with either pTMF3, pTMF3_Δ5UTR, or Fluc and HELZ2 followed by RT-qPCR (*i.e.*, using probe sets that specifically recognize the SV40 polyA signal of the plasmid, pTMF3, pTMF3_Δ5UTR, or Fluc RNAs) (**see Methods**) revealed that HELZ2 overexpression significantly reduced the RNA levels derived from the L1 5’UTR-containing constructs irrespective of their downstream sequences (**Fig. 6e**). Consistently, independent experiments in HeLa-JVM cells revealed that HELZ2 overexpression does not affect steady state RNA or protein levels produced from an inducible Tet-On firefly luciferase or human L1 ORFeus construct that lacks the L1 5’UTR^45^ (**Supplementary Figs. 6a and 6b**). Together, these data suggest that HELZ2 destabilizes L1 RNA by recognizing RNA sequences and/or RNA structure(s) within the L1 5’UTR.

Because previous experiments reported that L1 RNA induces a type I IFN response^40,42,46^ (**Fig. 2f**), we next tested whether the destabilization of L1 RNA by HELZ2 leads to a decrease in L1-mediated IFN-α induction. Strikingly, HELZ2 overexpression reduced the level of L1-dependent IFN-α induction to less than 5% of the control pJM101/L1.3FLAG construct (**Fig. 6f**, compare the leftmost and middle data graphs). Notably, this level of IFN-α induction was even lower than that observed in cells transfected with only the pCEP4 empty vector (**Fig. 6f**, compare the center and rightmost data graphs), raising the possibility that HELZ2 overexpression may also reduce the stablility of endogenous immunogenic RNAs to reduce basal levels of IFN-α induction.

## Discussion

Previous studies identified antiviral factors involved in innate immune responses that inhibit L1 retrotransposition by destabilizing L1 RNA, L1 proteins, L1 RNPs, and perhaps L1 (−) strand cDNAs (see Results: *“Proteins produced by Interferon-Stimulated Genes (ISGs) as potential L1 regulators”* for a complete list). In this study, we uncovered five additional ISG proteins that are enriched in IP/LC-MS/MS experiments conducted with WT ORF1p, but not the M8/RBM ORF1p mutant, which exhibits both attenuated RNA binding and L1 cytoplasmic foci formation.

HELZ2, HERC5, and OASL were predominantly localized in the cytoplasm, and upon overexpression, inhibited the retrotransposition of an engineered wild-type L1 (**Fig. 4a**; **Supplementary Fig. 4a**). Overexpression experiments further revealed that HELZ2 interacts with ORF1p in an RNA-dependent manner (**Figs. 3d and 6a**) and reduces the steady state levels of engineered L1 RNA, ORF1p, and ORF1p cytoplasmic foci formation (**Figs. 4b, 4c, 4d, and 4e**; **Supplementary Fig. 4b**). By comparison, HERC5 destabilizes ORF1p, but does not affect its cellular localization, whereas OASL mainly impairs ORF1p cytoplasmic foci formation. Thus, ISG proteins that predominantly act in the cytoplasm have the potential to inhibit L1 retrotransposition by acting at various steps in the L1 retrotransposition cycle (**Fig. 7**).

**Figure. 7:**
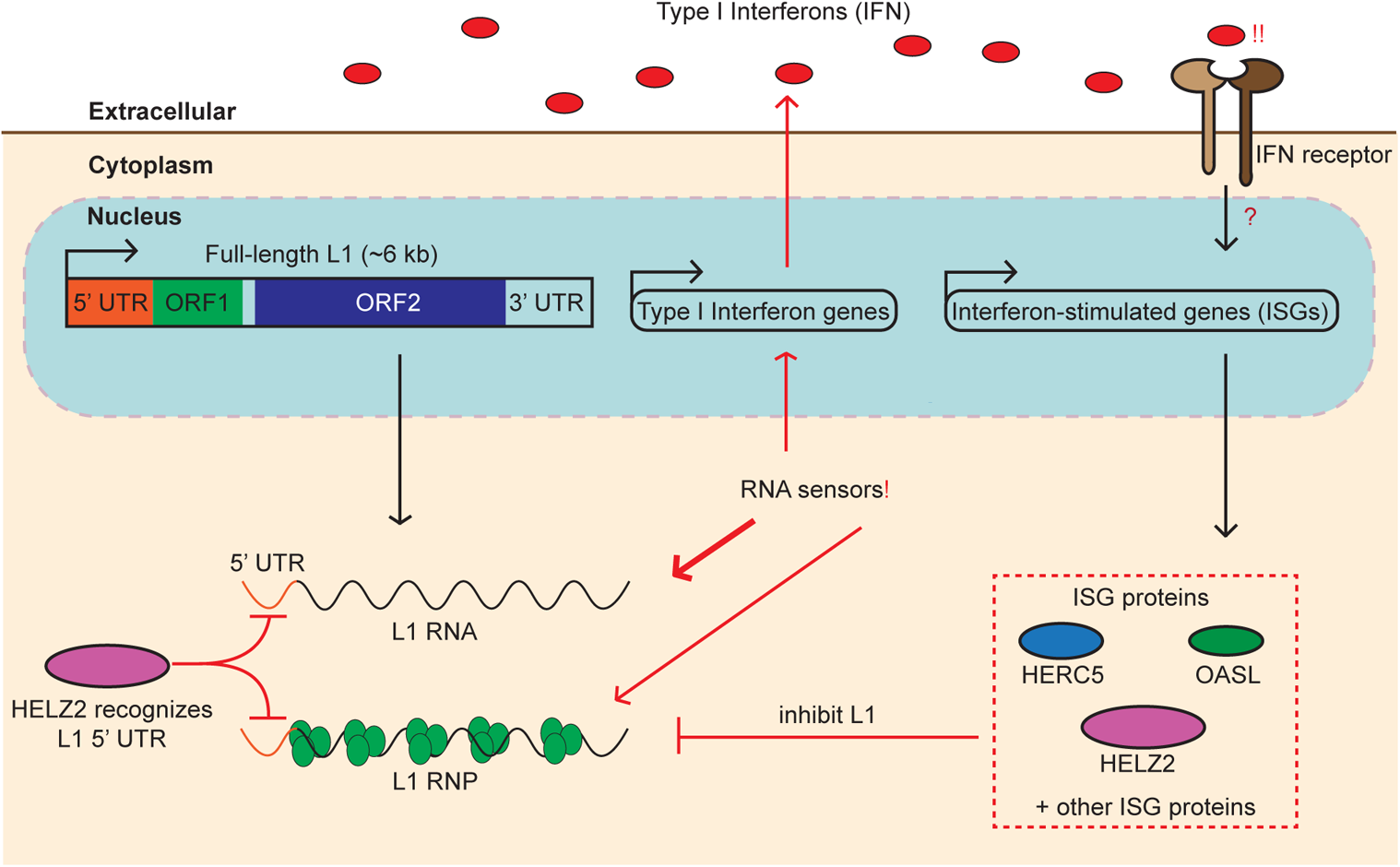
A working model hypothesizing a negative feedback loop between L1 RNA levels and ISG proteins. L1 RNAs and/or RNPs can be detected by cytoplasmic RNA sensors, which elicit the secretion of type I interferons (IFNs); ORF1p RNA-binding might shield L1 RNA from the sensors. IFN-binding to the extracellular IFN cell surface receptors then activates a signaling cascade, which induces the expression of ISGs, including *HELZ2*, *HERC5*, and *OASL*. These ISG proteins appear to inhibit L1 retrotransposition at different steps in the L1 retrotransposition cycle. HELZ2 appears to recognize RNA sequences and/or RNA structures within the L1 5’UTR, independently of ORF1p RNA binding, leading to the degradation of L1 RNA and subsequent blunting of the IFN response.

HELZ2 is a poorly characterized protein containing two putative RNA helicase domains that surround a centrally-located exoribonuclease (RNB) domain. RNB domains typically are flanked by cold shock and S1 RNA binding domains; however, HELZ2 appears to lack these domains^93^. A more in depth analysis of HELZ2 revealed similarities to other RNB-containing proteins, such as the prokaryotic cold shock inducible protein RNase R, which can degrade highly structured RNAs through its concerted helicase and 3’ to 5’ exoribonuclease activities^94,95^. Thus, it is tempting to suggest that HELZ2 might function in a similar stepwise manner, where its helicase activity initially unwinds L1 RNA secondary structures, allowing their subsequent degradation by the HELZ2 3’ to 5’ exoribonuclease activity (**Fig. 7**). Indeed, a HELZ2 helicase double mutant (WA1&2), but not a putative RNase-deficient mutant (dRNase), abolished the ability of HELZ2 to inhibit L1 retrotransposition (**Figs. 5b, 5c, 5d, and 5e**; **Supplementary Figs. 5c and 5d**), suggesting the HELZ2 helicase activity likely functions upstream of the single-strand 3’ to 5’ exoribonuclease activity to degrade L1 RNA. Because ORF1p-binding to L1 RNA is proposed to stabilize L1 RNAs^12,96^, we speculate that some regions of L1 RNA might be protected from HELZ2 degradation because of ORF1p RNA binding, but that other regions that have complex RNA secondary structures may be preferential HELZ2 targets. If so, HELZ2 might primarily destabilize these regions of L1 RNA to inhibit retrotransposition.

Previous studies demonstrated CpG DNA methylation of the L1 5’UTR is a potent means to inhibit endogenous L1 transcription^97–99^. DNA sequences within the 5’UTRs of older L1s (*e.g.*, members of the L1PA3 and L1PA4 subfamilies) also bind repressive Krüppel-associated Box-containing Zinc-Finger Protein 93 (ZNF93) to repress their transcription and deletion of these repressive sequences allowed the subsequent amplification of the L1PA2 and human-specific L1PA1 subfamilies in the human genome^100,101^. Notably, HELZ2 was discovered as a transcriptional co-activator of peroxisome proliferator-activated receptor alpha (PPAR-α)^102^ and PPAR-γ^103^. Because the L1 5’UTR also contains multiple transcription factor binding sites that drive L1 expression, it remains possible that HELZ2 might repress L1 transcription^21,22,104–106^.

Intriguingly, we found that L1 RNAs containing 5’UTR sequences are particularly susceptible to HELZ2-mediated RNA degradation (**Figs. 6e and 6f**; **Supplementary Fig. 6**), thereby representing a potential post-transcriptional mechanism by which RNA sequences and/or RNA structures within the 5’UTR are targeted by host proteins to inhibit L1 retrotransposition. Future studies are needed to test whether HELZ2 acts to destablilize the polypurine tract within the L1 3’UTR (which is absent from our expression vectors)^107^.

The overexpression of a WT L1 construct led to a modest upregulation of IFN-α expression, which previously was reported to contribute to inflammation, autoimmunity, and aging phenotypes^40,42–46^. A similar upregulation of IFN-α expression was observed upon the overexpression of an RT-deficient L1, and was slightly more pronounced upon the overexpression of the the ORF1p M8/RBM mutant. These data suggest that L1 RNA and or the L1-encoded proteins, but not intermediates generated during TPRT (*e.g.*, L1 cDNAs), are responsible for IFN-α upregulation in our experiments. Consistently, HELZ2 overexpression abolished L1-mediated IFN-α upregulation and also reduced IFN-α expression below baseline levels in our experiments, hinting that HELZ2 may also reduce the expression of endogenous immunogenic RNA(s).

Because L1s lack an extracellular phase in their replication cycle, one can posit that L1s would benefit from not triggering an innate immune response. That being stated, why the overexpression of the ORF1p M8/RBM mutant led to a more robust, yet modest, induction of IFN-α expression than the WT and RT-deficient L1s (**Fig. 2f**) requires further study. It is possible that ORF1p L1 RNA binding and/or the sequestration of L1 RNPs within cytoplasmic foci establishes effectively shields L1 RNAs from eliciting a interferon response and that the attenuated ability of the ORF1p M8/RBM mutant to bind L1 RNA could lead to higher levels of unprotected L1 RNA substrates that act as “triggers,” contributing to IFN-α expression (**Fig. 7**). Indeed, these data are consistent with a recent study, which reported that depletion of the Human Silencing Hub (HUSH complex) correlates with the derepression of primate-specific L1s and that the resultant L1 double-stranded RNAs may drive physiological or autoinflammatory responses in human cells^46^. Clearly, future studies are necessary to elucidate how and if L1 double-stranded RNAs, or perhaps single-stranded cDNAs, play important contributory roles to innate immune activation and human autoimmune diseases^108,109^.

## Materials and Methods

### Cell lines and cell culture conditions

The human HeLa-JVM cervical cancer-derived^17^, U-2 OS osteosarcoma-derived, and HEK293T embryonic carcinoma-derived cell lines were grown in Dulbecco’s Modified Eagle Medium (DMEM) (Nissui, Tokyo, Japan) supplemented with 10% fetal bovine serum (FBS) (Gibco, Amarillo, Texas, United States or Capricorn Scientific, Ebsdorfergrund, Germany), 0.165% NaHCO_3_, 100 U/mL penicillin G (Sigma-Aldrich, St. Louis, MO, United States), 100 µg/mL streptomycin (Sigma-Aldrich), and 2 mM L-glutamine (Sigma-Aldrich). HeLa-HA cells^69^ were grown in Minimum Essential Medium (MEM) (Gibco) supplemented with 10% FBS (Capricorn Scientific), 0.165% NaHCO_3_, 100 U/mL penicillin G, 100 µg/mL streptomycin, 2 mM L-glutamine, and 1x MEM Non-Essential Amino Acids Solution (Nacalai, Kyoto, Japan). The cell lines were grown at 37°C in 100% humidified incubators supplied with 5% CO_2_. The cell lines tested negative for mycoplasma contamination using a PCR-based method using the VenorGeM Classic Mycoplasma Detection Kit (Sigma-Aldrich). STR-genotyping was performed to confirm the identity of HeLa-JVM, HeLa-HA, U-2 OS, and HEK293T cells.

### Plasmid construction

#### Creation of the ORF1p-FLAG mutant constructs

Briefly, the pJM101/L1.3FLAG^7,50^ plasmid was used, unless otherwise indicated, to construct the plasmids in this study. Briefly, pJM101/L1.3FLAG DNA was used as a PCR template in conjunction with oligonucleotide primers containing the respective ORF1p mutations to generate the ORF1 mutants. The amplified PCR products and pJM101/L1.3FLAG plasmid DNA then were digested with *Not*I and *Age*I and ligated using the DNA Ligation Kit Mighty Mix (TaKaRa Bio, Shiga, Japan) at 16°C for 30 minutes. The resultant ligation products were transformed into *E*. *coli* XL1-Blue cells and plated on Luria Broth (LB) agar plates containing 50 µg/mL ampicillin. The resultant plasmids then were sequenced from the *Not*I to *Age*I restriction sites to ensure the integrity of the mutants.

#### Creation of the mCherry/GeBP1, ISG fusion protein, and HELZ2 mutant constructs

The *G3BP1* cDNA sequence was amplified from a HeLa-JVM total cDNA library and concurrently inserted in-frame with a *mCherry*-coding sequence into a lentiviral vector (pCW)^110^ using the in-Fusion Cloning Kit (TaKaRa Bio). The *HERC5*, *HELZ2*, *OASL*, *MOV10*, *IFIT1*, and *DDX60L* cDNAs were amplified from either a HeLa-JVM or HEK293T total cDNA library and inserted into the *Not*I and *Xho*I restriction sites in the pCMV-3Tag-9 vector (Agilent Technologies, Santa Clara, CA, United States) using either the Gibson Assembly Cloning Kit (New England Biolabs, Ipswich, MA, United States) or in-Fusion Cloning Kit. To generate *HELZ2* mutations, whole plasmid DNAs were amplified using primers harboring the intended mutations in separate reactions to avoid the formation of primer dimers. The template DNA then was digested with *Dpn*I at 37°C for 1 hour followed by heat inactivation at 80°C for 20 minutes. The PCR amplified DNA fragments were, mixed, annealed, and transformed into *E*. *coli* XL1-Blue cells.

### Plasmids used in this study

For mammalian cell experiments, plasmids were purified using the GenElute HP Plasmid Midiprep Kit (Sigma-Aldrich). All of the L1 expression plasmids contain a retrotransposition-competent L1 (L1.3, Genbank: L19088). The amino acid residues of ORF1p or ORF2p were counted from the first methionine of the L1.3 ORF1p and L1.3 ORF2p, respectively. The plasmids used in the study are listed below:

#### pCEP4 (Invitrogen)

the mammalian expression vector backbone used for cloning pJM101/L1.3 and pJJ101/L1.3 variants.

#### phrGFP-C (Agilent technology)

contains a humanized Renilla *GFP* gene whose expression is driven by a cytomegalovirus immediate-early (CMV) promoter.

#### pJM101/L1.3

was described previously^5,58^. This plasmid contains the full-length L1.3, cloned into the pCEP4 vector plasmid. L1 expression is driven by the CMV and L1.3 5’UTR promoters. The *mneoI* retrotransposition cassette was inserted into the L1.3 3’UTR as described previously^17^.

#### pJM101/L1.3FLAG

was described previously^50^. This plasmid is a derivative of pJM101/L1.3 that contain a single copy of the *FLAG* epitope tag fused in-frame to the 3’ end of the L1.3 *ORF1* sequence.

#### pJM105/L1.3

was described previously^25^. This plasmid is a derivative of pJM101/L1.3 that contains a D702A mutation in the ORF2p reverse transcriptase active site.

#### pTMF3

was described previously^92^. This plasmid is a derivative of pJM101/L1.3. A *T7 gene10* epitope tag was fused in-frame to the 3’ end of the *ORF1* sequence and three copies of a *FLAG* epitope tag were fused to the 3’ end of the *ORF2* sequence. This plasmid lacks the polypurine sequence in the L1 3’UTR.

#### pTMF3_Δ5UTR

is a derivative of pTMF3 that contains a deletion of the L1.3 5’UTR sequence.

#### pL1(5&3UTRs)_Fluc

is a derivative of pTMF3 that contains a firefly luciferase gene in place of the L1.3-coding region.

#### pJJ101/L1.3

was described previously^111^. This plasmid is similar to pJM101/L1.3, but contains an *mblastI* retrotransposition indicator cassette within the L1.3 3’UTR.

#### pJJ105/L1.3

was described previously^111^. This plasmid is a derivative of pJJ101/L1.3 that contains a D702A mutation in the ORF2p reverse transcriptase active site.

#### pALAF001_L1.3FLAG_M1

is a derivative of pJM101/L1.3FLAG that contains the N157A and R159A mutations in ORF1p, which abolished ORF1p cytoplasmic foci formation^48^.

#### pALAF002_L1.3FLAG_M2

is a derivative of pJM101/L1.3FLAG that contains the R117A and E122A mutations in ORF1p, which are proposed to adversely affect ORF1p trimerization^57^.

#### pALAF003_L1.3FLAG_M3

is a derivative of pJM101/L1.3FLAG that contains the N142A mutation in ORF1p, which is proposed to bind a chloride ion to stabilize ORF1p trimerization^12^.

#### pALAF004_L1.3FLAG_M4

is a derivative of pJM101/L1.3FLAG that contains the R135A mutation in ORF1p, which is proposed to bind a chloride ion to stabilize ORF1p trimerization^12^.

#### pALAF005_L1.3FLAG_M5

is a derivative of pJM101/L1.3FLAG that contains the E116A and D123A mutations in ORF1p, which are proposed to act as a binding site for host factors^12^.

#### pALAF006_L1.3FLAG_M6

is a derivative of pJM101/L1.3FLAG that contains the K137A and K140A mutations in ORF1p, which reduces the ability of ORF1p to bind L1 RNA^12^.

#### pALAF007_L1.3FLAG_M7

is a derivative of pJM101/L1.3FLAG that contains the R235A mutation in ORF1p, which reduces the ability of ORF1p to bind L1 RNA^49^.

#### pALAF008_L1.3FLAG_M8 (RBM)

is a derivative of pJM101/L1.3FLAG that contains the R206A, R210A, and R211A mutations in ORF1p, which severely impair the ability of ORF1p to bind L1 RNA^12^.

#### pALAF009_L1.3FLAG_M9

is a derivative of pJM101/L1.3FLAG that contains the R261A mutation in ORF1p, which reduces the ability of ORF1p to bind L1 RNA^49^.

#### pALAF010_L1.3FLAG_M10

is a derivative of pJM101/L1.3FLAG that contains the Y282A mutation in ORF1p, which is proposed to reduce nucleic chaperone activity^49^.

#### pALAF012_mCherry-G3BP1_pCW

contains the *mCherry* sequence fused in frame to a human *G3BP1* cDNA in a lentiviral expression vector, pCW^110^. The puromycin resistant gene and reverse tetracycline-controlled trans-activator (rtTA) coding regions are in-frame and are expressed by a human PGK promoter; puromycin and rtTA are separated by a self-cleaving T2A peptide so that each protein can be expressed from the bicistronic transcript. The *mCherry-G3BP1* cDNA is expressed from a doxycycline inducible (Tet-On) promoter. In the presence of doxycycline, rtTA can adopt an altered confirmation that allows it to bind the Tet-On promoter to allow *mCherry-G3BP1* expression.

#### pALAF015_hHELZ2L-3xMYC

contains the canonical human *HELZ2* long isoform cDNA (2649 bps) cloned into pCMV-3Tag-9 (Agilent Technologies), which allows the expression of a HELZ2-3xMYC fusion protein. The CMV promoter drives *HELZ2-3xMYC* expression.

#### pALAF016_hIFIT1-3xMYC

contains the human *IFIT1* cDNA cloned into pCMV-3Tag-9, which allows the expression of a hIFIT1-3xMYC fusion protein. The CMV promoter drives *IFIT1-3xMYC* expression.

#### pALAF021_hDDX60L-3xMYC

contains the human *DDX60L* cDNA cloned into pCMV-3Tag-9, which allows the expression of a hDDX60L-3xMYC fusion protein. The CMV promoter drives *DDX60L-3xMYC* expression.

#### pALAF022_hOASL-3xMYC

contains the human *OASL* cDNA cloned into pCMV-3Tag-9, which allows the expression of the OASL-3xMYC fusion protein. The CMV promoter drives *OASL-3xMYC* expression.

#### pALAF023_hHERC5-3xMYC

contains the human *HERC5* cDNA cloned into pCMV-3Tag-9, which allows the expression of a HERC5-3xMYC fusion protein. The CMV promoter drives *HERC5-3xMYC* expression.

#### pALAF024_hMOV10-3xMYC

contains the human *MOV10* cDNA cloned into pCMV-3Tag-9, which allows the expression of a MOV10-3xMYC fusion protein. The CMV promoter drives *MOV10-3xMYC* expression.

#### cepB-gfp-L1.3

was described previously^92^. The plasmid contains the full-length L1.3 with an *EGFP* retrotransposition reporter cassette, *mEGFPI*. L1.3 expression is augmented by the L1 5’UTR promoter. The plasmid backbone also contains a *blasticidin S-deaminase* (*BSD*) selectable marker driven by the SV40 early promoter.

#### cepB-gfp-L1.3RT(−)

was described previously^92^. The plasmid is identical to cepB-gfp-L1.3 but contains a D702A mutation in the ORF2p reverse transcriptase active site.

#### cepB-gfp-L1.3RT(−) intronless

was described previously^92^. The plasmid is similar to cepB-gfp-L1.3RT(−) except that the intron in the *mEGFPI* retrotransposition cassette was removed, allowing EGFP expression in the absence of L1.3 retrotransposition.

#### cep99-gfp-L1.3

was described previously^92^. The plasmid is similar to cepB-gfp-L1.3 but contains the puromycin resistant gene instead of the blasticidin resistance gene as a selectable marker.

#### cep99-gfp-L1.3RT(−) intronless

was described previously^92^. The plasmid is similar to cep99-gfp-L1.3 except that it contains the D702A mutation in the ORF2p reverse transcriptase domain and the intron in the *mEGFPI* retrotransposition cassette was removed, allowing EGFP expression in the absence of L1.3 retrotransposition.

#### pALAF025_hHELZ2L-3xMYC_WA1

is a derivative of pALAF015_hHELZ2L-3xMYC that contains the K550A mutation in the Walker A motif of the N-terminal HELZ2 helicase domain, which is predicted to inactivate the ATP binding ability of the helicase domain^88^.

#### pALAF026_hHELZ2L-3xMYC_WA2

is a derivative of pALAF015_hHELZ2L-3xMYC that contains the K2180A mutation in the Walker A motif of the carboxyl-terminal HELZ2 helicase domain, which is predicted to inactivate the ATP binding ability of the helicase domain^88^.

#### pALAF027_hHELZ2L-3xMYC_WA1&2

is a derivative of pALAF015_hHELZ2L-3xMYC that contains the K550A and K2180A mutations in the Walker A motifs of both HELZ2 helicase domains^88^.

#### pALAF028_hHELZ2L-3xMYC_WB1

is a derivative of pALAF015_hHELZ2L-3xMYC that contains the E668A mutation in the Walker B motif of the N-terminal helicase domain of HELZ2, which is predicted to inactivate the ATP hydrolysis activity of the helicase domain^88^.

#### pALAF029_hHELZ2L-3xMYC_WB2

is a derivative of pALAF015_hHELZ2L-3xMYC that contains the E2361A mutation in the Walker B motif of the C-terminal helicase domain of HELZ2, which is predicted to inactivate the ATP hydrolysis activity of the helicase domain^88^.

#### pALAF030_hHELZ2L-3xMYC_dRNase

is a derivative of pALAF015_hHELZ2L-3xMYC that contains the D1346N, D1354N, and D1355N mutations in the RNB domain of HELZ2, which is predicted to inactivate the RNase activity of the RNB domain^87^.

#### psPAX2

is a lentivirus packaging vector that was a gift from Didier Trono (Addgene plasmid # 12260). The plasmid expresses the HIV-1 gag and pol proteins.

#### pMD2.G

is a lentivirus envelope expression vector that was a gift from Didier Trono (Addgene plasmid # 12259). The plasmid expresses a viral envelope protein and the vesicular stomatitis virus G glycoprotein (VSV-G).

#### pcDNA6

was described previously^92^. It is a derivative of pcDNA6/TR (Invitrogen, Carlsbad, CA, United States) and contains the *blasticidin S-deaminase* (*BSD*) selectable marker but lacks the TetR gene. This plasmid was made by Dr. John B. Moldovan (University of Michigan Medical School).

#### pCMV-3Tag-8-Barr

is a human *β*-*Arrestin* expression plasmid. The human *ARRB2* cDNA was cloned into pCMV-3Tag-8 (Agilent Technologies). The plasmid contains three copies of a *FLAG* epitope tag fused in-frame to the 3’ end of the *ARRB2* cDNA. The CMV promoter drives *ARRB2-3xFLAG* expression.

#### pTMO2F3_Alu

is plasmid that co-expresses Alu and a monocistronic version of L1 ORF2p that contains the L1 5’UTR. The monocistronic *ORF2* coding sequence contains three copies of an in-frame *FLAG* epitope tag sequence at its 3’ end; the CMV promoter augments the expression of *ORF2-3xFLAG*. The plasmid also contains an AluY element whose expression is driven by a 7SL promoter. The Alu element contains the *neo*^Tet^ retrotransposition indicator cassette^91^, which was inserted upstream of the Alu poly(dA) tract. This arrangement allows the quantification of Alu retrotransposition efficiency by counting the resultant number of G418-resistant foci. This plasmid lacks the polypurine sequence in the L1 3’UTR.

#### pTMO2F3D145AD702A_Alu

is identical to pTMO2F3_Alu but contains the D145A and D702A mutations, which inactivate the ORF2p endonuclease and reverse transcriptase activities, respectively.

#### pTMO2H3_Alu

is a derivative of pTMO2F3_Alu plasmid where the *FLAG* epitope tag was replaced with three copies of *HA* epitope tag sequence.

#### pSBtet-RN

was a gift from Eric Kowarz^45,112^ (Addgene plasmid # 60503). The plasmid contains a firefly luciferase (*Fluc*) gene with an upstream Tet-On inducible promoter.

#### pDA093

was a gift from Kathleen Burns^45^ (Addgene plasmid # 131390). This plasmid is similar to pSBtet-RN but the luciferase gene was replaced with the human L1 *ORFeus* (*ORF1* and *ORF2*) sequence lacking the 5’ or 3’UTR.

#### pCMV(CAT)T7-SB100

was a gift from Zsuzsanna Izsvak^113^ (Addgene plasmid # 34879). This plasmid contains a hyperactive variant of the *Sleeping Beauty* transposase, whose expression is driven by the CMV promoter.

### Western blots

HeLa-JVM, U-2 OS, or HEK293T cells were seeded in a 6-well tissue culture plate (Greiner, Frickenhausen, Germany) at 2×10^5^ cells per well. On the following day, the cells were transfected with 1 μg of DNA (1 μg of an L1-expressing plasmid or 0.5 μg of the L1-expressing plasmid and 0.5 μg of either a pCMV-3Tag-8-Barr control or ISG-expressing plasmid) using 3 µL of FuGENE HD transfection reagent (Promega, Madison, WI, United States) and 100 µL of Opti-MEM (Gibco) according to the protocol provided by the manufacturer. The medium was replaced with fresh DMEM approximately 24 hours post-transfection (day 1). The cells were harvested using 0.25% trypsin (Gibco) at days 2 through 9 post-transfection (depending on the specific experiment). The transfected cells were enriched using 100 µg/mL of hygromycin B (Wako, Osaka, Japan), which was added to the media two days post-transfection and replaced with fresh DMEM containing hygromycin B daily. After collection by trypsinization, the cells were pelleted by centrifugation at 300 × *g* for 5 minutes. Then, the cells were washed twice with cold 1x PBS, flash-frozen in liquid nitrogen, and kept at −80°C.

For cell lysis, the cells were incubated in Radio-ImmunoPrecipitation Assay (RIPA) buffer (10 mM Tris-HCl [pH 7.5], 1 mM EDTA, 1% TritonX-100, 0.1% sodium deoxycholate, 0.1% SDS, 140 mM NaCl, 1x cOmplete EDTA-free protease inhibitor cocktail [Roche, Mannheim, Germany]) at 4°C for 30 minutes. The cell debris was pelleted at 12,000 × *g* for 5 minutes and the supernatant was collected. The protein concentration was measured using the Protein Assay Dye Reagent Concentrate (Bio-Rad, Richmond, CA, United States) and all of the samples for each experiment were normalized to the same concentration. The protein lysate was mixed at an equal volume with 3x SDS sample buffer (187.5 mM Tris-HCl [pH 6.8], 30% glycerol, 6% SDS, 0.3M DTT, 0.01% bromophenol blue) and boiled at 105°C for 5 minutes. Twenty micrograms of total protein lysates for all samples were separated using sodium dodecyl sulphate-polyacrylamide gel electrophoresis (SDS-PAGE). Proteins on the gel were transferred onto a Immobilon-P, 0.45 μm pore, polyvinylidene difluoride (PVDF) transfer membrane (Merck Millipore, Billerica, MA, United States) using 10 mM CAPS buffer (3-[cyclohexylamino]-1-propanesulfonic acid [pH 11]) in a Mini Trans-Blot Electrophoretic Transfer Cell tank (Bio-Rad) according to protocol provided by the manufacturer. The transfer was performed at 4°C at 50V for 16 hours. After the transfer was completed, the membrane was incubated with Tris-NaCl-Tween (TNT) buffer (0.1 M Tris-HCl [pH 7.5], 150 mM NaCl, 0.1% Tween 20) containing 3% skim milk (Nacalai) for 30 minutes. The membranes then were washed with TNT buffer, cut into strips, and incubated with the relevant primary antibodies in TNT buffer at 4°C overnight. The next day, the membranes were washed four times with TNT buffer with five minutes interval at room temperature and incubated with HRP-conjugated secondary antibodies in TNT buffer containing 0.01% SDS at room temperature for an hour. The membranes were washed four times with TNT buffer with five minutes interval at room temperature and the signals were detected with the Chemi-Lumi One L (Nacalai) chemiluminescence reagent using a LAS-3000 Imager (Fujifilm, Tokyo, Japan), LAS-4000 Imager (Fujifilm), or a FUSION Solo S Imager (Vilber-Lourmat, Marne-la-Vallee, France).

#### Primary antibodies and dilutions (in parentheses)

Mouse monoclonal anti-FLAG M2 antibody (1/5000), (Sigma-Aldrich, F1804, RRID: AB_262044)

Mouse monoclonal anti-MYC antibody (1/5000), (Cell Signaling Technology, 9B11, RRID: AB_331783)

Rabbit polyclonal anti-PABPC1 antibody (1/5000), (Abcam, ab21060, RRID: AB_777008)

Mouse monoclonal anti-GAPDH antibody (1/5000), (Millipore, MAB374, RRID: AB_2107445) Mouse monoclonal anti-Actin antibody (1/5000), (Millipore, MAB1501, RRID: AB_2223041) Rabbit polyclonal anti-T7-tag antibody (1/5000), (Cell Signaling Technology, D9E1X, RRID: AB_2798161)

Goat polyclonal anti-Luciferase antibody (1/2000), (Promega, G7451, RRID: AB_430862) Mouse monoclonal anti-ORF1p (4H1) antibody (1/2000), (Millipore, MABC1152)

Mouse monoclonal anti-eIF3 p110 (B-6) antibody (1/5000), (Santa Cruz Biotechnology, sc-74507, RRID: AB_1122487)

#### Secondary antibodies and dilutions (in parentheses)

Sheep polyclonal anti-mouse HRP-conjugated Whole antibody (1/5000), (GE Healthcare, NA931-1ML, RRID: AB_772210)

Goat polyclonal anti-rabbit HRP-conjugated Whole antibody (1/5000), (Cell Signaling Technology, 7074, RRID: AB_2099233)

Donkey polyclonal anti-rabbit HRP-conjugated Whole antibody (1/5000), (GE Healthcare, NA934-1ML, RRID: AB_772206)

Donkey polyclonal anti-goat HRP-conjugated Whole antibody (1/5000), (Santa Cruz Biotechnology, sc-2020, RRID: AB_631728)

### Immunofluorescence

#### Cell transfection and fixation

HeLa-JVM or U-2 OS cells were plated on 18 mm glass coverslips (Matsunami Glass, Osaka, Japan) coated with Alcian Blue 8GX (Sigma-Aldrich) in 12-well tissue culture plates (Greiner) at 2.5×10^4^ cells per well in DMEM (with 1.0 μg/mL of doxycycline in mCherry-G3BP1-expressing U-2 OS cells). After 24 hours, the cells were transfected with 0.5 µg of plasmid DNA (0.5 µg of the L1-expressing plasmid [pJM101/L1.3FLAG, pALAF002, pALAF005, or pALAF008] or 0.25 µg of pJM101/L1.3FLAG and 0.25 µg of either a pCMV-3Tag-8-Barr control or ISG-expression plasmid) using 1.5 µL of FuGENE HD transfection reagent and 50 µL of Opti-MEM according to protocol provided by the manufacturer. Approximately 24 hours post-transfection, the medium was replaced with fresh DMEM and 1.0 μg/mL of doxycycline was added into the medium for mCherry-G3BP1-expressing U-2 OS cells. Approximately 48 hours post-transfection, the cells were washed with 1x PBS and fixed with 4% paraformaldehyde (PFA) at room temperature for 15 minutes. Prior to cell fixation, the cells were treated with DMSO (Sigma-Aldrich) or 0.5 mM sodium meta-arsenite (Sigma-Aldrich) for one hour. The fixed cells then were washed with 1x PBS three times and kept at 4°C until cell permeabilization.

#### Immunostaining

The resultant cells were permeabilized with 0.2% Triton X-100 and 0.5% normal donkey serum (NDS) for 5 minutes. The cells were washed once with 1x PBS and twice with PBST (1x PBS and 0.1% Tween 20) following permeabilization. The primary antibodies (1/1000 dilution in PBST) containing 0.5% NDS were applied onto the coverslip and incubated for 45 minutes at room temperature. The cells were washed with PBST three times after the primary antibody incubation. The secondary antibodies (1/250 dilution in PBST) containing 0.5% NDS and 0.1 μg/mL of 4’, 6-diamidino-2-phenylindole (DAPI) were applied onto the coverslip and incubated for 45 minutes at room temperature. The cells were washed with PBST three times followed by multiple rinses with water. The excess liquid was removed, and the glass coverslips were fixed on glass slides with 3 µL of VECTASHIELD (Vector Laboratories, Burlingame, CA, United States).

#### Immunofluorescence

Images were captured using the DeltaVision Elite microscope (Cytiva, Marlborough, MA, United States). Six z-stack images with 1 µm thickness difference were captured and projected into a single image with the max intensity for each image. For ORF1p-FLAG probed with the Alexa 488-conjugated antibody or MYC-tagged proteins probed with the Cy5-conjugated antibody, the FITC/AF488 or Cy5/AF647 channel was used, respectively. mCherry-G3BP1 fluorescence was detected through the AF594/mCherry channel. In the ORF1p foci counting experiments, the same signal intensity threshold was applied to all samples and only cells with visible ORF1p signals were counted as positive cells. Only cells that displayed clear cytoplasmic ORF1p signals with foci distinguishable from the background were counted as an L1 foci-positive cells.

#### Primary antibodies and dilutions (in parentheses)

Mouse monoclonal anti-FLAG M2 antibody (1/1000), (Sigma-Aldrich, F3165, RRID: AB_259529)

Rabbit polyclonal anti-FLAG antibody (1/1000), (Sigma-Aldrich, F7425, RRID: AB_439687) Mouse monoclonal anti-MYC antibody (1/1000), (Cell Signaling Technology, 9B11, RRID: AB_331783)

#### Secondary antibodies and dilutions (in parentheses)

Donkey anti-mouse polyclonal Alexa Fluor 488 IgG (H+L) (1/250), (Thermo Fisher Scientific, A-21202, RRID: AB_141607)

Donkey anti-rabbit polyclonal Alexa Fluor 488 IgG (H+L) (1/250), (Thermo Fisher Scientific, A-21206, RRID: AB_2535792)

Goat polyclonal anti-mouse Cy5 (1/250), (Jackson ImmunoResearch Labs, 115-175-146, RRID: AB_2338713)

### Lentiviral transduction

HEK293FT cells were plated in a 10-cm tissue culture dish at 1×10^6^ cells per plate. On the following day, the cells were transfected with 5 µg plasmid DNA (2.5 µg of pALAF012, 1.875 µg of psPAX2, and 0.625 µg of pMD2.G) using 15 µL of 1 mg/mL transfection grade linear polyethylenimine hydrochloride (MW 40,000) (PEI-MAX-40K) (Polysciences, Warrington, PA, United States) in 500 µL of Opti-MEM. Approximately 24 hours post-transfection, the medium was replaced with fresh DMEM. The medium containing the virus was collected 48 hours post-transfection and filtered through a 0.45 µm polyethersulfone (PES) filter (Merck Millipore, Billerica, MA).

To generate the inducible mCherry-G3BP1-expressing U-2 OS cell line, 2×10^5^ cells per well were plated in a 6-well tissue culture plate. On the next day, the medium was replaced with virus-containing medium supplemented with 8 µg/mL of polybrene (Sigma-Aldrich). Approximately 24 hours post-viral treatment, the medium was replaced with fresh DMEM. From the second-day post-viral treatment onwards, the media was replaced with fresh DMEM containing 1 µg/mL puromycin every three days until the non-transduced cells were dead.

### Construction of cell lines expressing Tet-On Luciferase and human L1 ORFeus

HeLa-JVM cells were plated in 6-well plates at 2×10^5^ cells per well. On the following day, the cells were transfected with 500 ng of plasmid DNA (pSBtet-RN or pDA093) and 50 ng of a sleeping beauty plasmid (pCMV[CAT]T7-SB100) using 2.0 µL of FuGENE HD transfection reagent and 100 µL of Opti-MEM according to the protocol provided by the manufactures. After ~24 hours, the medium was replaced with fresh DMEM. G418 (Nacalai) selection (500 µg/mL) began ~48 hours post-transfection for 1 week; the G418 containing media was replaced daily. Five percent of the total living cells were transferred into 10-cm tissue culture dishes and the media was replaced daily with 500 µg/mL G418 until the cells reached ~90% confluency. The cells then were trypsinized and resuspended in PBS containing 2% FBS and dTomato-positive cells were sorted using a BD FACSAria III flow cytometer (BD Biosciences, San Jose, CA, United States) to obtain clonal cell lines. Western blotting was used to screen the resultant cell lines for doxycycline dosage-dependent expression of Luciferase or human L1 ORFeus.

### L1 and Alu Retrotransposition Assays

L1 or Alu cultured cell retrotransposition assays were performed as described with modifications^17,54,58,59,91,114^.

In retrotransposition assays using the *mneoI* retrotransposition indicator cassette, 2×10^5^ HeLa-JVM or HeLa-HA cells per well were seeded in 6-well tissue culture plates. On the following day, the cells were transfected with 1 µg of DNA (0.5 µg of pJM101L1.3/FLAG or its variants and 0.5 µg of phrGFP-C for the L1 retrotransposition assay) or 1 µg of DNA (0.5 µg of pTMO2F3_Alu or phrGFP-C and 0.5 µg of pCMV-3Tag-8-Barr control, pALAF015 [HELZ2], or pALAF024 [MOV10] for the Alu retrotransposition assay) using 3 µL FuGENE HD and 100 µL of Opti-MEM according to the protocol provided by the manufacturer. The medium was replaced with fresh DMEM (HeLa-JVM) or MEM (HeLa-HA), respectively ~24 hours post-transfection (day 1). On day 3 post-transfection, to check transfection efficiency, each duplicate was collected, fixed with 0.5% paraformaldehyde, and subjected to flow cytometry analysis using BD Accuri C6 Plus Flow Cytometer (BD Biosciences). The FITC channel was used to determine the number of hrGFP-expressing cells out of 10, 000 cells as a transfection efficiency control. The medium in the remaining transfectants was replaced daily with fresh DMEM or MEM containing 500 µg/mL G418 from day 3 onwards. The resultant colonies were fixed at day 10-14 post-transfection using the fixation solution (1x PBS containing 0.2% glutaraldehyde and 2% formaldehyde). The cells were stained with 0.1% crystal violet. The resultant number of foci were counted and normalized to the transfection efficiency. Please note: the HEK293T cells are G418-resistant and could not be used in *mneoI* based retrotransposition assays.

In retrotransposition assays using the *mblastI* retrotransposition indicator cassette, 5×10^4^ HeLa-JVM cells per well were seeded in 6-well tissue culture plates. After ~24 hours, the cells were transfected with 1 µg of DNA (0.5 µg of pJJ101/L1.3 and 0.5 µg of an ISG-expressing plasmid or pCMV-3Tag-8-Barr) using 3 µL of FuGENE HD in 100 µL of Opti-MEM. For the viability control, 5×10^3^ HeLa-JVM cells per well were seeded in 6-well tissue culture plates. After ~24 hours, the cells were transfected with 1 µg of DNA (0.5 µg of pcDNA6 and 0.5 µg of an ISG-expressing plasmid or pCMV-3Tag-8-Barr) using 3 µL of FuGENE HD in 100 µL of Opti-MEM. Approximately 24 hours post-transfection (day 1), the medium was changed with fresh DMEM. Blasticidin selection (10 µg/mL of blasticidin S HCl) began from day 4 post-transfection and the media containing blasticidin was replaced every three days until day 8-10. The resultant colonies were fixed using the fixation solution and stained with 0.1% crystal violet. The resultant number of foci were counted and normalized to the resultant number of pcDNA6-transfected foci.

In retrotransposition assays using the *mEGFPI* retrotransposition indicator cassette, 2×10^5^ HeLa-JVM or HEK293T cells per well were seeded in 6-well tissue culture plates. On the next day, the cells were transfected with 1 µg of DNA (0.5 µg of cepB-gfp-L1.3 or cepB-gfp-L1.3RT[-] intronless and 0.5 µg of a pCMV-3Tag-8-Barr control or ISG-expressing plasmid) using 3 µL of FuGENE HD in 100 µL of Opti-MEM. Approximately 24 hours post-transfection (day 1), the medium was replaced with fresh DMEM. Transfected cells were selected using 10 µg/mL blasticidin S HCl from day 2 post-transfection, changing the media every three days. The cells were collected on day 7-8 post-transfection and the resultant EGFP positive cells were analyzed using BD Accuri C6 Plus Flow Cytometer. The FITC channel was used to count the EGFP positive cells out of 30,000 cells. The number of the EGFP-positive cells was normalized to the transfection efficiency measured by counting the number of cepB-gfp-L1.3RT(−) intronless GFP-positive cells.

### siRNA treatment

HeLa-JVM cells were plated in 6-well tissue culture plates at 1×10^5^ cells per well. After ~24 hours, 25 nM of a Dharmacon siRNA mixture (non-targeting control: ON-TARGETplus Non-targeting Pool, D-001810-10-0020; HELZ2: ON-TARGETplus HELZ2 siRNA SMARTpool, L-019109-00-0005; or MOV10: ON-TARGETplus MOV10 siRNA SMARTpool, L-014162-00-0005) were transfected using 3.75 μL of Lipofectamine RNAiMAX (Thermo Fisher Scientific, Waltham, MA, United States). Approximately 24 hours post-siRNA treatment (day 1), the medium was replaced with fresh DMEM and the cells were transfected with 0.5 µg of cepB-gfp-L1.3 or cepB-gfp-L1.3RT(−) intronless using 1.5 μL of FuGENE HD in 100 μL of Opti-MEM. Transfected cells were selected using 10 μg/mL blasticidin S HCl from day 3 post-transfection with media changes every three days. On day 8 post-transfection, the cells were harvested, washed with cold 1x PBS twice, and analyzed for EGFP expression using BD Accuri C6 Plus Flow Cytometer out of 30,000 cells. The number of the EGFP-positive cells was normalized to the transfection efficiency measured by counting the number of cepB-gfp-L1.3RT(−) intronless GFP-positive cells.

### Immunoprecipitation of L1 ORF1p

#### Immunoprecipitation for IP-MS

HeLa-JVM cells were plated in 15-cm tissue culture dishes containing DMEM medium at 1.5×10^6^ cells per dish. Three 15-cm tissue culture dishes were used for each sample preparation. After ~24 hours, the cells were transfected with 15 μg of an L1-expressing plasmid (pJM101/L1.3, pJM101/L1.3FLAG or pALAF008) using 45 μL of FuGENE HD (Promega) in 1,500 μL of Opti-MEM. On the following day (day 1), the medium was replaced with fresh DMEM. From day 2 post-transfection onwards, the medium was replaced daily with fresh DMEM containing 100 μg/ml hygromycin B. On day 6 post-transfection, the cells were harvested using trypsin, washed with 1x cold PBS twice, flash-frozen with liquid nitrogen, and stored at −80°C.

For IP reactions, one hundred fifty microliters of Dynabeads Protein G (Invitrogen) was washed twice with PBS containing 0.5% BSA and 0.1% Triton X-100. For each sample, the beads were incubated with 15 μg of mouse monoclonal anti-FLAG M2 antibody (Sigma-Aldrich, F1804, RRID: <u>AB_262044</u>) in 1 mL of PBS containing 0.5% BSA and 0.1% Triton X-100 at 4°C for 2 hours. After incubation, the antibody-conjugated beads were washed with PBS containing 0.5% BSA and 0.1% Triton X-100 twice. The beads were resuspended in Lysis150 buffer (20 mM Tris-HCl [pH 7.5], 2.5 mM MgCl_2_, 150 mM KCl, 0.5% IGEPAL CA-630, 1 mM DTT) containing 0.2 mM phenylmethylsulfonyl fluoride (PMSF) and 1x cOmplete EDTA-free protease inhibitor cocktail before immunoprecipitation. Each cell pellet was lysed using the Lysis150 buffer containing 0.2 mM PMSF and 1x cOmplete EDTA-free protease inhibitor cocktail. The resuspended cell pellets were incubated at 4°C for 30 minutes and centrifuged at 12,000 × *g* for 5 minutes to pellet the cell debris. The supernatant was collected and incubated with antibody non-conjugated Dynabeads Protein G at 4°C for 2 hours with gentle rotation to remove non-specific protein binding. The Dynabeads were removed and the protein concentration in the pre-cleared cell lysates was quantified using Protein Assay Dye Reagent Concentrate. The same total amount of protein was used for each immunoprecipitation. Dynabeads Protein G conjugated to the anti-FLAG antibody was added to the supernatant and incubated at 4°C for 3 hours with gentle rotation. The beads were then washed five times with 200 μL of the Lysis150 buffer. The ORF1p-FLAG protein complex bound was eluted using 200 μg/mL of 3xFLAG peptide (Sigma-Aldrich) in the Lysis150 buffer containing 0.2 mM PMSF and 1x cOmplete EDTA-free protease inhibitor cocktail by incubation at 4°C for 1 hour with gentle rotation. This step was repeated once and the protein was precipitated overnight using cold acetone. The protein was pelleted at 12,000 × *g* at 4°C for 30 minutes, resuspended in 1x SDS sample buffer and boiled at 105°C for 5 minutes.

#### Immunoprecipitation for western blotting

HEK293T cells were plated in 10-cm tissue culture dishes at 3×10^6^ cells per dish. Approximately 24 hours after plating, the cells were transfected with 4 μg of pJM101/L1.3FLAG or pJM101/L1.3 and 2 μg of ISG-expressing plasmid (pALAF015, pALAF016, pALAF021, pALAF022, pALAF023, or pALAF024) using 18 μL of 1 mg/mL PEI-MAX-40K in 500 μL of Opti-MEM. Approximately 24 hours post-transfection, the media was changed with fresh DMEM. From day 2 post-transfection onwards, the medium was replaced daily with fresh DMEM containing 100 μg/ml hygromycin B. On day 4 post-transfection, the cells were harvested with pipetting, washed with 1x cold PBS twice, flash-frozen with liquid nitrogen, and stored at −80°C for subsequent experiments.

For each sample, ten microliters of the Dynabeads Protein G were incubated with 1 μg of anti-FLAG M2 antibody in 50 μL of PBS containing 0.5% BSA and 0.1% Triton X-100 at 4°C for 2 hours. After incubation, the antibody-conjugated beads were washed with PBS containing 0.5% BSA and 0.1% Triton X-100 twice. The beads were resuspended in Lysis150 buffer containing 0.2 mM PMSF and 1x cOmplete EDTA-free protease inhibitor cocktail before immunoprecipitation. Each cell pellet was lysed in 500 μL of the Lysis150 buffer containing 0.2 mM PMSF and 1x cOmplete EDTA-free protease inhibitor cocktail. The resuspended cell pellets were incubated at 4°C for 1 hour and centrifuged at 12,000 × *g* for 5 minutes to pellet the cell debris. The supernatant was collected and 10 μL of the supernatant was saved as input. Anti-FLAG antibody-conjugated Dynabeads were added to the samples and incubated at 4°C for 4 hours with gentle rotation.

The RNase treatment for HELZ2-expressed samples was performed after removal of the cell lysate using 20 μg/mL of RNase A (Nippongene, Tokyo, Japan) in 100 μL of the Lysis150 buffer for five minutes at 37°C. The beads then were washed four times with 100 μL of the

Lysis150 buffer. The beads were resuspended directly in 1x SDS sample buffer and boiled at 105°C for 5 minutes except for the HELZ2-expressed samples, where the ORF1p-FLAG protein complex was eluted using 20 μL of the Lysis150 buffer containing 0.2 mM PMSF, 1x cOmplete EDTA-free protease inhibitor cocktail, and 200 μg/mL 3xFLAG peptide by incubation at 4°C for 1 hour with gentle rotation. The eluted protein was resuspended in 1x SDS sample buffer and boiled at 105°C for 5 minutes.

### Proteomic analysis by LC-MS/MS

Mass spectrometry analysis was performed by the proteomics facility in the Graduate School of Biostudies at Kyoto University. After SDS-PAGE and visualization of the gel using PlusOne Silver Staining Kit, Protein (Cytiva) according to the protocol provided by the manufacturer, the entire gel lane from each sample was excised into 15 components. The silver stain was then removed, and the excised gel slices were incubated with sequencing-grade modified trypsin (Promega) to extract the peptides. The purified peptides then were subjected to liquid chromatography-tandem mass spectrometry (LC-MS/MS) on nano-Advance (AMR, Tokyo, Japan) and Q Exactive Plus (Thermo Fisher Scientific). The MS/MS spectra and protein scores were analyzed using the Proteome Discoverer 1.4 (Thermo Fisher Scientific) and the MASCOT server 2.5.1 (Matrix Science: https://www.matrixscience.com/) against UniProt Knowledgebase (UniProtKB: https://www.uniprot.org/help/uniprotkb). Keratin proteins were removed from all the lists. To identify the ORF1p-FLAG interacting proteins, the peptide matches for each UniProt accession number obtained from the WT L1 (pJM101/L1.3FLAG)-expressing cells were compared to that of the no tag control (pJM101/L1.3) or the RBM L1 (pALAF008)-expressing cells.

### ORF1p crystal structure analysis

The crystal structure images of ORF1p and the mutations were created using UCSF ChimeraX software 1.2.5 for Windows^115^ based on the 2ykp pdb file^12^.

### GO term analysis

Protein hits obtained from IP-MS with five peptide matches or more were submitted for Gene Ontology (GO) analysis using the PANTHER statistical enrichment analysis tool^55^ (http://pantherdb.org). The UniProt accession ID with the respective number of peptide matches was submitted for GO biological process complete annotation data set analysis. The FDR value was used for the correction.

### GSEA Leading Edge Analysis

GSEA 4.1.0 for Windows software was used for the analysis^56^ (http://www.broad.mit.edu/GSEA) using the hallmark gene sets from GSEA Molecular Signatures Database (<u>MSigDB:</u> https://www.gsea-msigdb.org/gsea/msigdb/). The leading edge analysis was performed on the GSEA results using the GSEA 4.1.0 software. Peptide matches of WT ORF1p-FLAG *vs*. M8/RBM-FLAG were compared and the enriched gene sets were subjected to leading edge analysis. All of the UniProt accession ID hits (1255 Uniprot accession numbers) from IP-MS with the respective number of peptide matches were included in the analysis.

#### ImageJ quantification of western blot band intensity

Using the ImageJ software tool^116^, identical sized rectangles were drawn for each band. The area of intensity of the bands were generated using Plot Lanes function and calculated using a wand (tracing) tool. The intensity of each ORF1p-T7 band was normalized to that of the GAPDH band with respective samples. The values were displayed as ratios in comparison to the leftmost band in the western blot image (pTMF3 and pCMV-3Tag-8-Barr control co-transfected cells).

### RNA extraction and RT-qPCR

HeLa-JVM or HEK293T at 2×10^5^ cells per well were seeded in 6-well tissue culture plates. On the following day, the cells were transfected with 1 μg of DNA (1 μg of an L1-expressing plasmid or 0.5 μg of the L1-expressing plasmid and 0.5 μg of a pCMV-3Tag-8-Barr control or an ISG-expressing plasmid). Approximately 24 hours post-transfection (day 1), the medium was replaced with fresh DMEM. On day 2 (HeLa-JVM and HeLa-HA) or day 4 (HEK293T) post-transfection, the cells were washed with 1x PBS and 0.9 mL TRIzol was added directly to each well. The RNA extractions were performed according to the protocol provided by the manufacturer. The cells were lysed with TRIzol and transferred into new 1.5 mL tubes. One hundred eighty microliters of chloroform was added into each tube and shaken vigorously for 15 seconds. After incubation at room temperature for 5 minutes, the samples were centrifuged at 12,000 × *g* for 15 minutes at 4°C. Three hundred sixty microliters of the upper layer were transferred into a new 1.5 mL tube and 400 μL of 100% isopropanol was added to precipitate the RNA. The samples were incubated at room temperature for 10 minutes. Next, RNA was pelleted at 12,000 × *g* for 30 minutes. The purified RNA then was washed with 75% cold ethanol and centrifuged at 10,000 × *g* for 5 minutes. The RNA pellet was dried at room temperature. Once dried, 30 μL of RNase-free H_2_O was added and incubated at 55°C for 10 minutes to dissolve RNA. The resultant RNA was then treated with RNase-free DNase Set (QIAGEN) according to the protocol provided by the manufacturer with some minor modifications. Five microliters of DNase I (15 K units, TaKaRa Bio), 0.2 U/μL of ribonuclease inhibitor (porcine liver) (TaKaRa Bio) in 44.5 μL of the RNase-free Buffer RDD was added to each sample. The samples were incubated at room temperature for 15 minutes and the RNA then was pelleted after ethanol precipitation (incubation at −20°C overnight in 240 μL of 100% ethanol and 8 μL of 3M NaOAc [pH 5.2]). The RNA pellets were washed with 75% cold ethanol, dried at room temperature, resuspended in RNase-free water, and incubated at 75°C for 10 minutes to inactivate the DNase I. One microgram of total RNA was used as a template in reverse transcription reactions using 0.2 mM dNTP (TakaRa Bio), 1 U/μL ribonuclease inhibitor (porcine liver) (TaKaRa Bio), 0.25 U/μL AMV reverse transcriptase XL (TaKaRa Bio), and 0.125 μM of an oligo (dT) primer (Invitrogen) according to the protocol provided by the manufacturer unless stated otherwise. Two negative controls were included for all instances: no reverse transcriptase (reverse transcriptase was excluded during cDNA synthesis) and no template (cDNA was replaced with RNase-free water). The reverse transcription reaction was performed as follows: 30°C for 10 minutes, 42°C for 30 minutes, and 95°C for 5 minutes. Prime Script MMLV reverse transcriptase (TaKaRa Bio) and 0.125 μM of the oligo (dT) primer for RNA-IP experiments (see below) or a HELZ2 specific primer (HELZ2_R) for HELZ2 RNA quantification were used to reverse transcribe instead. RNA was incubated at 65°C for 5 minutes before the addition of Prime Script MMLV reverse transcriptase and the reverse transcription was performed as follows: 42°C for 60 minutes followed by 70°C for 15 minutes. RT-qPCR was performed using Luna Universal qPCR Master Mix (New England Biolabs). Amplification was performed using StepOnePlus Real-Time PCR System (Applied Biosystems) using the following parameters: 15 seconds at 95°C; followed by 40 cycles of denaturation (95°C for 15 seconds) and amplification (60°C for 60 seconds). Technical duplicates were made for each sample. Quantification of cDNA for each reaction was determined by comparing the cycle threshold (Ct) with a standard curve generated from one of the samples using StepOne Software v2.2. All Ct readings fall within the range of the standard curve generated.

#### Primers used for RT-qPCR

L1 (SV40)_F: 5’-TCCAGACATGATAAGATACATTGATGAG-3’

L1 (SV40)_R: 5’-GCAATAGCATCACAAATTTCACAAA-3’

L1 (FLAG)_F: 5’-ATGGATTACAAGGACGACGATG-3’

L1 (FLAG)_R: 5’-TGTGTGAATTTGATCCTGTCAT-3’

Luciferase_F: 5’-CGAGGCTACAAACGCTCTCA-3’

Luciferase_R: 5’-CAGGATGCTCTCCAGTTCGG-3’

IFN-α _F: 5’-CTGAATGACTTGGAAGCCTG-3’

IFN-α _R: 5’-ATTTCTGCTCTGACAACCTC-3’

HELZ2_F: 5’-GAGAAGGTGGTTCTTCTCGGAG-3’

HELZ2_R: 5’-CTCATGCATGCGGTACTGAG-3’

MOV10_F: 5’-CGTACCGGAAACAGGTGGAG-3’

MOV10_R: 5’-TGAACCCACCTTCAAGTCCTTG-3’

mneoI (Alu or L1)_F: 5’-ACCGGACAGGTCGGTCTTG-3’

mneoI (Alu or L1)_R: 5’-CTGGGCACAACAGACAATCG-3’

Beta-actin_F: 5’-CCTTTTTTGTCCCCCAACTTG-3’

Beta-actin_R: 5’-TGGCTGCCTCCACCCA-3’

GAPDH_F: 5’-GGAGTCCCTGCCACACTCAG-3’

GAPDH_R: 5’-GGTCTACATGGCAACTGTGAGG-3’

Oligo (dT): 5’-TTTTTTTTTTTTTTTTTTTTVN-3’

### RNA-IP

RNA immunoprecipitation (RNA-IP) experiments were carried out as described previously with some modifications^27^. HeLa-JVM cells were plated in 10-cm tissue culture dishes at 1.5×10^6^ cells per dish. On the following day (day 0), the cells were transfected with 5 μg of plasmid DNA (pJM101/L1.3, pJM101/L1.3FLAG or pALAF008_M8) using 15 μL of PEI-MAX-40K in 500 μL of Opti-MEM. Approximately 24 hours post-transfection (day 1), the medium was replaced with fresh DMEM. On the following day (day 2), the medium was replaced daily with fresh DMEM containing 100 μg/mL hygromycin B and the cells were collected at day 5 post-transfection. The whole cell extracts were prepared by incubation in the Lysis150 buffer containing 0.2 mM PMSF and 1x cOmplete EDTA-free protease inhibitor cocktail for one hour at 4°C. The lysate was separated from the insoluble fraction by centrifugation at 12, 000 × *g* for five minutes and transferred to a new tube. Ten microliters of the lysate were saved as the input fraction. Prior to immunoprecipitation, the anti-FLAG antibody-conjugated beads were prepared as described in “immunoprecipitation and western blotting” section of the Methods. The cleared lysate (input) was incubated with the anti-FLAG antibody-conjugated beads for 5 hours at 4°C. The beads were then washed four times with 150 μL of Lysis150 buffer without protease inhibitors. The RNA extraction was performed as described in “RNA extraction and RT-qPCR” in the Methods section with a slight modification: 200 μg/mL glycogen was added to the immunoprecipitated RNA fraction before ethanol precipitation. All of the RNA samples were resuspended in 30 μL of RNase-free water. Five microliters (one sixth) of the extracted RNA from the input and IP fractions were used to synthesize cDNA using PrimeScript MMLV reverse transcriptase as described in the previous section. The ORF1p-associated RNA values were calculated by dividing the cDNA amount in the IP fraction by that in the input.

### Statistical analysis

One-way ANOVA followed by Bonferroni-Holm post hoc tests were performed for all statistical analyses unless stated otherwise. All analyses were performed using online website statistical calculator ASTATSA (https://www.astatsa.com/) or GraphPad Prism version 9.0.0 for Windows (GraphPad Software, San Diego, California USA; www.graphpad.com). The numbers of biological replicates are indicated in the figure legends. Data are shown as the mean ± standard errors of the means (SEM). The *p*-value of each pair was indicated in the figure legends. ns: not significant; * *p<*0.05; ** *p*<0.01; *** *p<*0.001).

### Data availability

IP-MS data are available in the source data file.

## Acknowledgements

We thank K. H. Burns, D. Ardeljan, T. Heidmann, M. T. Hayashi, D. Trono, and Z. Izsvak for valuable reagents, Ishikawa lab members, T. Makino, K. Sugino, S. Matsuo, K. Onishi, K. Nishimori, S. Adachi, K. Yamaguchi, M. Miyoshi, and Dr. J. B. Moldovan for helpful discussions. A.L-F. was supported by JASSO and MEXT Scholarships. F.I. was supported by JSPS KAKENHI (Grant Number JP19H05655). J.V.M. was supported by NIH grant GM060518. T.M. was supported by JSPS KAKENHI (Grant Number JP18K06180 and 21K19219), ISHIZUE 2021 of Kyoto University Research Development Programs, and research grants from the Takeda Science Foundation, the Japan Foundation for Applied Enzymology, the Sumitomo Foundation for Basic Science Research Projects, and Astellas Foundation for Research on Metabolic Disorders.

## Author information

Affiliations

**Department of Gene Mechanisms, Graduate School of Biostudies, Kyoto University, Kyoto 606-8501, Japan**

Ahmad Luqman-Fatah, Fuyuki Ishikawa, Tomoichiro Miyoshi

**Proteomics Facility, Graduate School of Biostudies, Kyoto University, Kyoto 606-8501, Japan**

Yuzo Watanabe

**Radiation Biology Center, Graduate School of Biostudies, Kyoto University, Kyoto 606-8501, Japan**

Ahmad Luqman-Fatah, Fuyuki Ishikawa, Tomoichiro Miyoshi

**Department of Human Genetics, University of Michigan Medical School, Ann Arbor, MI 48109-5618, USA**

John V. Moran

**Department of Internal Medicine, University of Michigan Medical School, Ann Arbor, MI 48109-5618, USA**

John V. Moran

## Contributions

A.L-F., J.V.M., and T.M. conceived, designed, analyzed data and prepared the manuscript. A.L-F. and T.M. performed experiments. Y.W. provided technical support and performed mass spectrometry. F.I., J.V.M., and T.M. contributed with critical discussion, reading and editing. All authors contributed to ideas.

## Corresponding author

Please address correspondence to Tomoichiro Miyoshi.

## Ethics declaration

Competing interests

J.V.M. is an inventor on patent US6150160, is a paid concultant for Gilead Sciences, serves on the scientific advisory board to Tessera Therapeutics Inc. (where he is paid as a consultant and has equity options), has licensed reagents to Merck Pharmaceutical, and recently served on the the American Society of Human Genetics Board of Directors. The other authors declare no competing interests.

## Supplementary information

Supplementary Figures 1-6

Supplementary Tables 1-3

## Source data

Source data file

---

**Supplementary Figure. 1 (supporting Fig. 1a and Supplementary Figs. 2a and 2b):**
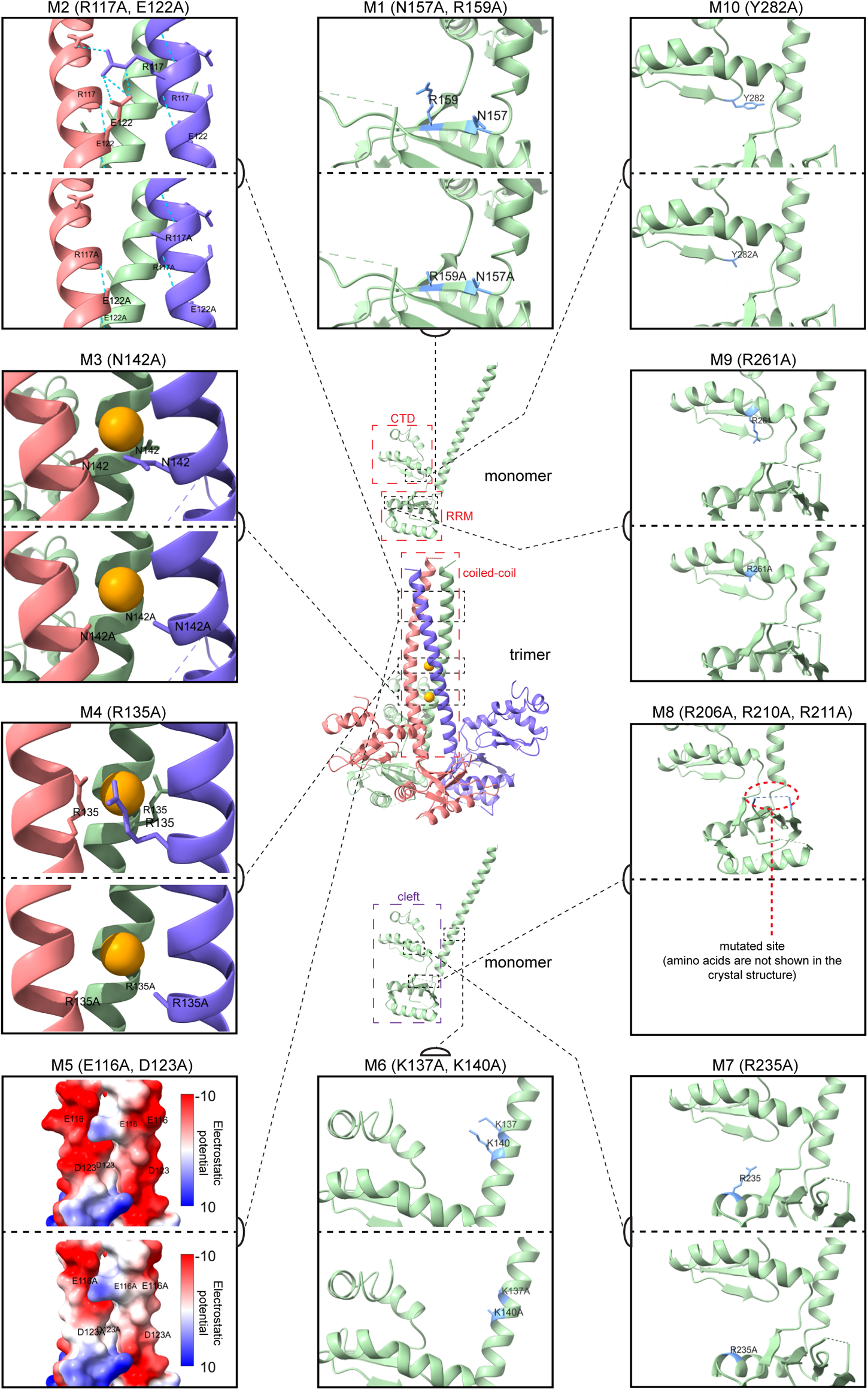
Crystal structure of L1 ORF1p mutants. **Center:** *Crystal structure of the ORF1p trimer (middle) and monomer (top and bottom)*. Shown is the crystal structure assembly of the ORF1p trimer from amino acid residues 107 to 323 in the “lifted” conformation (Protein Data Bank ID: 2ykp); each monomer is annotated with distinct colors (green, purple, and red colors). Two chloride ion residues (orange spheres) are shown in the predicted position inside the coiled-coil domain (red-dotted box, top of the trimer). Each monomer forms a flexible cleft (indicated in bottom monomer: purple-dotted box) made up of an RNA recognition motif (top monomer: RRM, bottom of the cleft, red-dotted box) and a C-terminal domain (top monomer: CTD, top of the cleft, red-dotted box) to bind RNA. Relative positions of the mutated amino acids are indicated in black-dotted boxes connected with black-dotted lines to the respective enlarged images of the mutated sites. **Periphery:** *Mutated sites of the ORF1p mutants in this study*. Based on the number from lowest (*e.g.,* M1 and M2) to highest (*e.g.*, M10), the ORF1p mutants were arranged in a counterclockwise direction beginning from the top (middle), where each of the mutant is enclosed in black boxes with the respective annotation noted at the top. Amino acids corresponding to the WT and alanine missense mutations are indicated within each box, where the WT (upper) and the mutants (lower) are separated by black-dotted lines. M1, M6, M7, M9 and M10: mutated amino acids and side chains are indicated in blue. M2: blue-dotted lines indicate hydrogen bonds formed, including between R117 and E122 side chains (different monomers) to stabilize the trimer. M3 and M4: depicted are the predicted side chains thought to stabilize the chloride ions. M5: a relative electrostatic potential map of the ORF1p trimers surface, the mutated site was suggested to be a potential recruitment site of host factors. Red indicates low positive electrostatic potential (high acidity) and blue indicates high positive electrostatic potential (high basicity). M8: the mutated site is shown in the red-dotted circle.

**Supplementary Figure. 2 (supporting Figs. 1a, 1b, and 1d):**
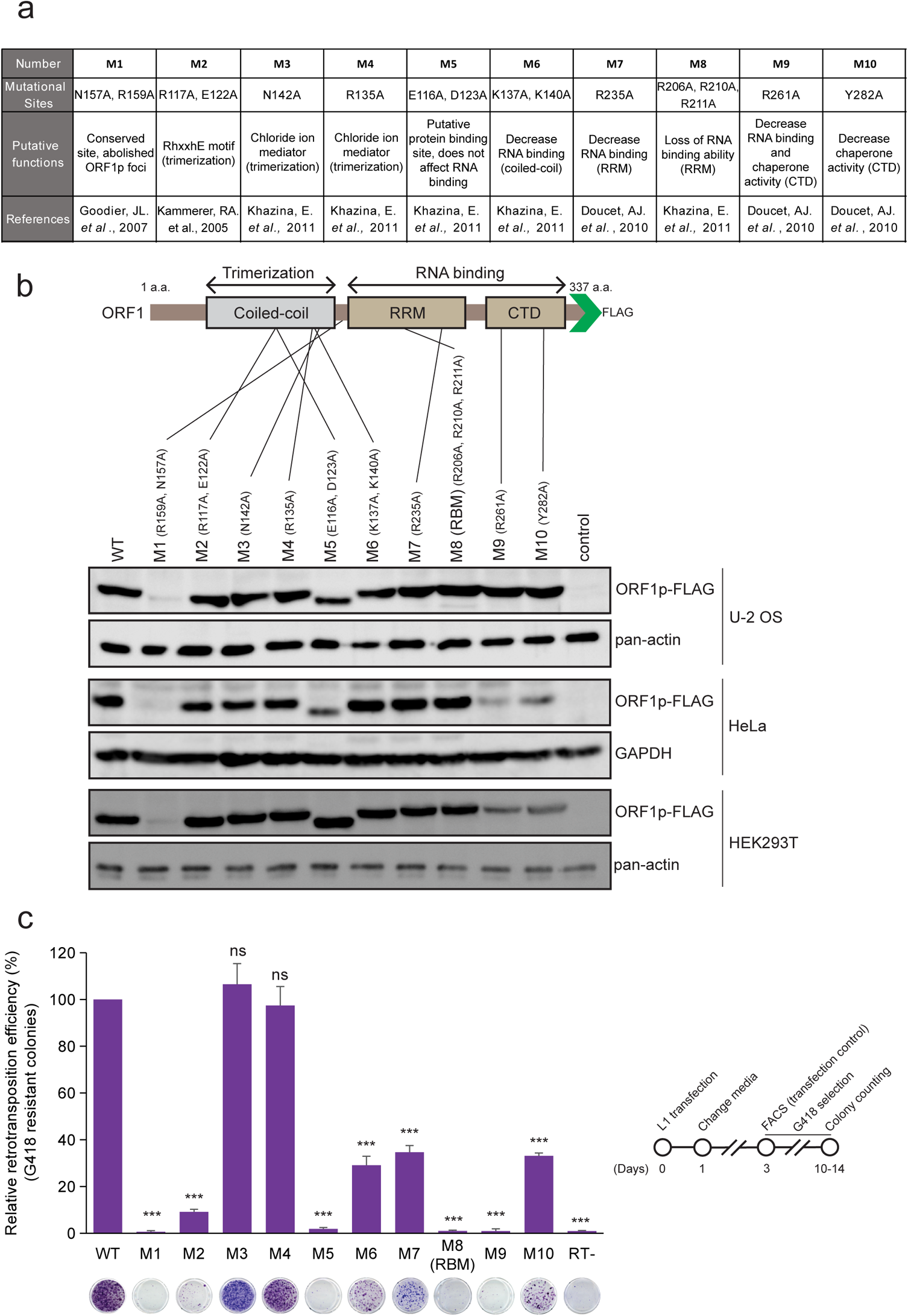
L1 ORF1p mutational analyses. **(a)** *ORF1p mutants generated in this study.* Ten alanine missense ORF1p-FLAG mutants (M1 to M10) were tested in various assays. Row 1, mutant number. Row 2, alanine mutations; commas denote double (*i.e.*, M1, M2, M5, M6) or triple (*i.e.*, M8) mutants. Row 3, putative functional domains affected by the alanine mutations. Row 4, references to previous studies implicating the mutations in L1 biology. Some of the mutants were designed based upon the ORF1p crystal structure. **(b)** *Schematic representation of ORF1p functional domains containing the mutations noted in panel (a).* Top, relative positions of the respective mutated amino acids. Bottom, western blots to test whether the relative mutations are expressed in U-2 OS, HeLa-JVM, or HEK293T cells. The cells were transfected with: pJM101/L1.3FLAG (WT); pALAF001 (M1); pALAF002 (M2); pALAF003 (M3); pALAF004 (M4); pALAF005 (M5); pALAF006 (M6); pALAF007 (M7); pALAF008 (M8); pALAF009 (M9); or pALAF010 (M10). U-2 OS, HeLa-JVM, or HEK293T cells were collected on day 5, day 9, or day 4 post-transfection, respectively. An anti-FLAG antibody was used to detect ORF1p-FLAG. Pan-actin and GAPDH served as loading controls. **(c)** *L1 retrotransposition efficiencies.* HeLa-JVM cells were co-transfected with the plasmids used in panel (b) and a phrGFP-C plasmid to normalize for transfection efficiencies and subjected to *mneoI*-based retrotransposition assays (inset, timeline of the assay). X-axis, mutant name and representative results from the assay; a missense mutation in the ORF2p RT domain (RT-) served as a negative control. Y-axis, the percentage of normalized G418-resistant foci compared to the WT (pJM101/L1.3FLAG) control. Pairwise comparisons relative to the WT control: *p* = 1.8 × 10^−12^*** (M1); 7.6 × 10^−12^*** (M2); 0.56^ns^ (M3); 0.67^ns^ (M4); 2.1 × 10^−12^*** (M5); 5.7 × 10^−10^*** (M6); 1.4 × 10^−9^*** (M7); 2.1 × 10^−12^*** (M8); 2.0 × 10^−12^*** (M9); 1.3 × 10^−9^*** (M10); 2.0 × 10^−12^*** (RT-). Values represent the mean ± SEM of three independent biological replicates. The *p*-values were calculated using a one-way ANOVA followed by Bonferroni-Holm post-hoc tests: ns: not significant; *** *p<*0.001. The relative retrotransposition efficiencies and the representative results reported in Fig. 1d were taken from this retrotransposition assay (WT and M8 [RBM]).

**Supplementary Figure. 3 (supporting Figs. 1e and 1f):**
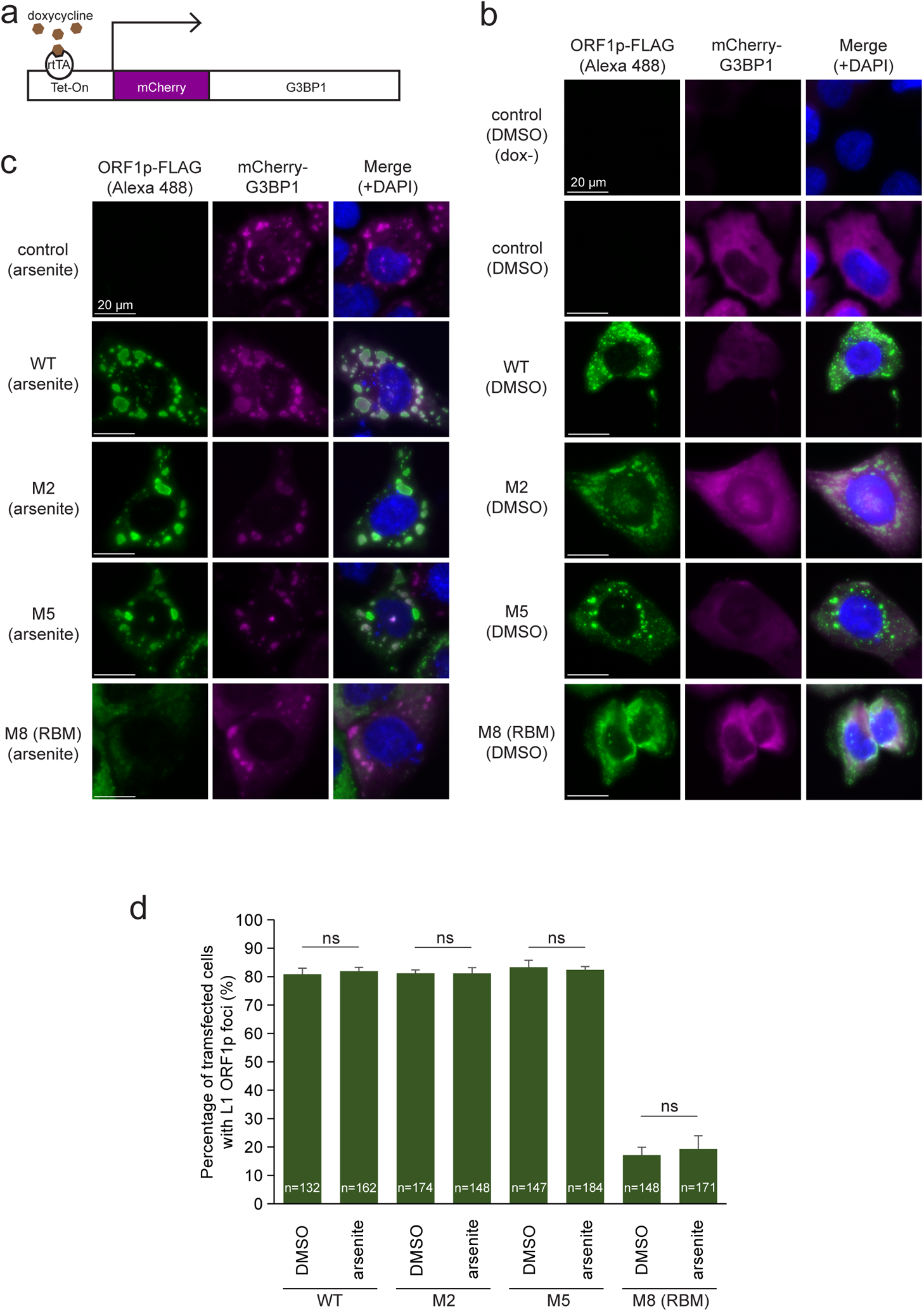
L1 cytoplasmic foci formation with the ORF1p mutants. **(a)** *Schematic of the doxycycline inducible mCherry-G3BP1 expression plasmid*. An mCherry-G3BP1 fusion protein only will be expressed in U-2 OS cells when doxycycline binds to the reverse tetracycline-controlled *trans*-activator protein (rtTA) and rtTA subsequently binds to the Tet-On promoter to activate mCherry-G3BP1 transcription. **(b and c)** Representative immunofluorescence images of WT, M2, M5, and M8 (RBM) ORF1p localization in the absence (b) or presence (c) of arsenite. U-2 OS cells containing the inducible mCherry-G3BP1 expression cassette were transfected with pCEP4 (control), pJM101/L1.3FLAG (WT), pALAF002 (M2), pALAF005 (M5), or pALAF008 (M8). Two days post-transfection, the cells were treated with DMSO or 0.5 mM sodium arsenite for 1 hour prior to fixation. A mouse primary anti-FLAG antibody and secondary anti-mouse-Alexa Fluor 488 fluorescent dye-conjugated antibodies were used to visualize ORF1p. Cells not treated with doxycycline (dox-) were included as a control in panel (b). White bars, 20 µm. **(d)** Quantification of ORF1p-FLAG cytoplasmic foci in U-2 OS cells transfected with WT, M2, M5, or M8 (RBM) ORF1p L1 expression constructs. X-axis, construct name and whether the cells were treated with vehicle (DMSO) or arsenite. Y-axis, the percentage of transfected cells exhibiting ORF1p-FLAG cytoplasmic foci. The numbers (n) within the green rectangles indicate the number of cells analyzed in the experiment. The percentage of transfected cells with L1 ORF1p foci data in Fig. 1f were taken from the WT (DMSO) and M8 (RBM) (DMSO) samples. Pairwise comparisons between DMSO and arsenite-treated cells: *p* = 1.00^ns^ (WT); 1.00^ns^ (M2); 1.00^ns^ (M5); 1.00^ns^ (M8 [RBM]). Values represent the mean ± SEM of three independent biological replicates. The *p*-values were calculated using a one-way ANOVA followed by Bonferroni-Holm post-hoc tests. ns: not significant.

**Supplementary Figure. 4 (supporting Figs. 4a and 4b):**
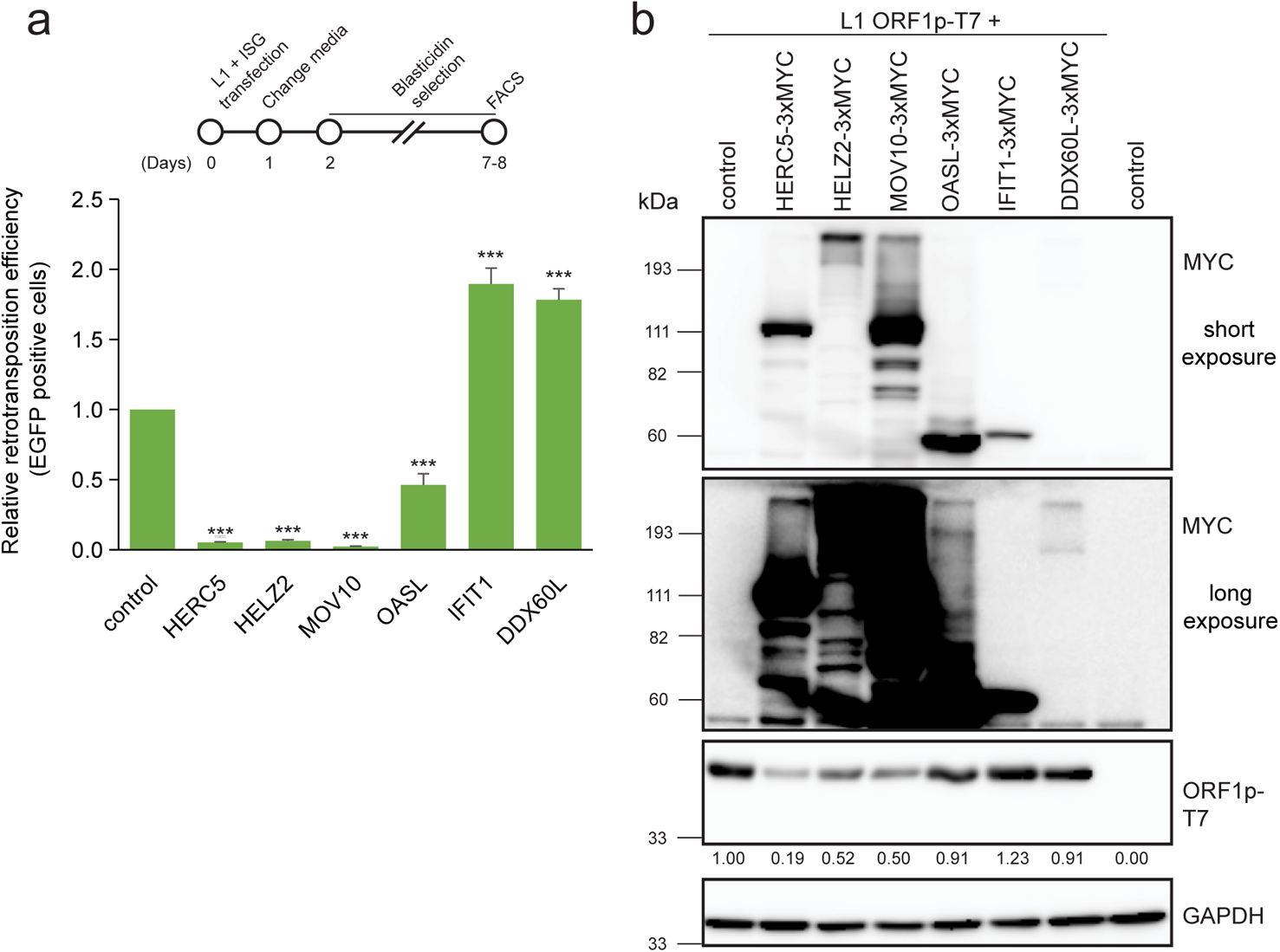
Functional analysis of the ISG proteins in HEK293T cells. **(a)** *Overexpression of HERC5, HELZ2, and OASL inhibit L1 retrotransposition in HEK293T.* Top: the timeline of the assay. HEK293T cells we co-transfected with cep99-gfp-L1.3 (which has the *mEGFPI* retrotransposition indicator cassette) and either pCEP4 (control) or the following individual ISG protein expression plasmids containing three copies of a MYC epitope tag (3xMYC) at their respective carboxyl termini: pALAF015 (HELZ2); pALAF016 (IFIT1); pALAF021 (DDX60L); pALAF022 (OASL); pALAF023 (HERC5); or pALAF024 (MOV10). EGFP-positive cells transfected with cep99-gfp-L1.3 were counted using flow cytometry and normalized to the number of EGFP-positive cells in the transfection control (*i.e.*, cells independently transfected with the cep99-gfp-L1.3RT(−) intronless plasmid and each of the above listed plasmids). X-axis, name of constructs co-transfected with cep99-gfp-L1.3. Y-axis, relative percentage of EGFP-positive cells relative to the cep99-gfp-L1.3 + pCEP4 control. Pairwise comparisons relative to the control: *p* = 4.8 × 10^−7^*** (HERC5); 4.6 × 10^−7^*** (HELZ2); 6.1 × 10^−7^*** (MOV10); 3.9 × 10^−5^*** (OASL); 6.2 × 10^−7^*** (IFIT1); 1.5 × 10^−6^*** (DDX60L). Values represent the mean ± SEM from three independent biological replicates. The *p*-values were calculated using a one-way ANOVA followed by Bonferroni-Holm post-hoc tests (*** *p<*0.001). **(b)** *Western blot detection of ORF1p in HEK293T cells co-transfected with ISG-expressing plasmids*. HEK293T cells were co-transfected with pTMF3 (L1 containing T7 epitope-tagged ORF1p) and either pCMV-3Tag-8-Barr (control) or the individual ISG-expressing plasmids used in panel (a). The relative band intensities of ORF1p-T7 are indicated under the ORF1p-T7 blot. They were calculated using ImageJ software and are normalized to the respective GAPDH band intensities. An anti-MYC antibody was used to detect the ISG proteins, and the western blot was shown as the short (top) and long exposure (bottom) images. An anti-T7 antibody was used to detect WT ORF1p-T7. GAPDH served as a loading control. Molecular weight markers (kDa) are indicated at the left of the blots.

**Supplementary Figure. 5 (supporting Figs. 5b, 5c, 5d, and 5e):**
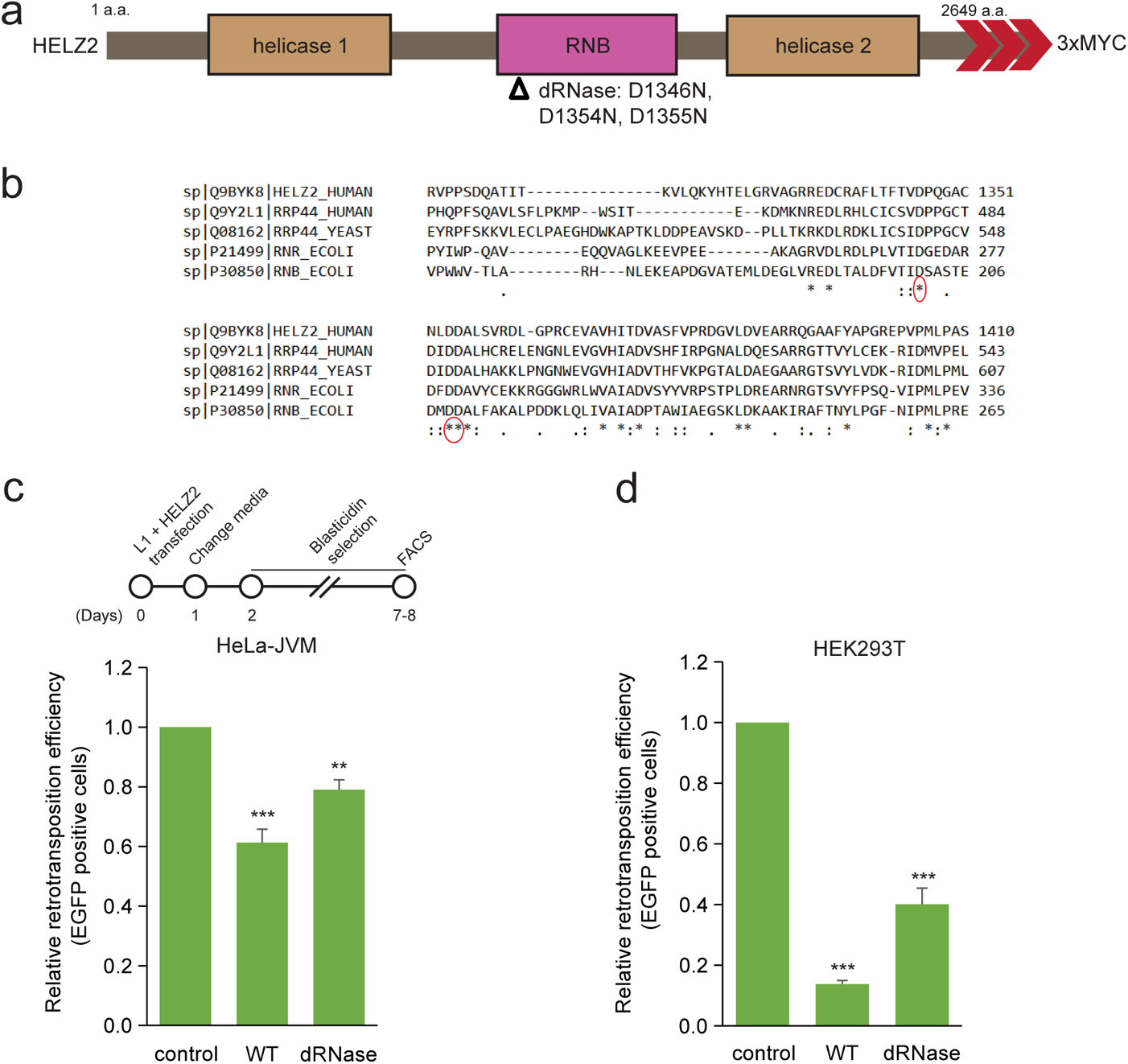
Functional analyses of the HELZ2 RNB domain. **(a)** *Schematic of mutations in the HELZ2 RNB domain.* The HELZ2 protein contains two putative helicase domains (helicase 1 and helicase 2), which surround a putative RNB exonuclease domain. Open triangle, position of the missense mutations in conserved amino acids within the RNB domain: D1346N/D1354N/D1355N (dRNase). Three red arrowheads, relative positions of the 3xMYC carboxyl-terminal epitope tags. **(b)** *Identification of conserved amino acids in the RNB domain.* Multiple sequence alignments of the following RNB-containing proteins: *Homo sapiens* exosome complex exonuclease Rrp44 (RRP44_HUMAN) and HELZ2 (HELZ2_HUMAN); *Saccharomyces cerevisiae* exosome complex exonuclease Rrp44 (RRP44_YEAST); and *Escherichia coli* RNase R (RNR_ECOLI) and Exoribonuclease 2 (RNB_ECOLI). Red circles, amino acids mutated in the D1346N/D1354N/D1355N (dRNase) triple mutant. **(c)** *L1 retrotransposition efficiency in the presence of the D1346N/D1354N/D1355N (dRNase) mutant in HeLa-JVM cells*. Top: the timeline for the retrotransposition assays shown in panels (c) and (d). HeLa-JVM cells were co-transfected with cepB-gfp-L1.3 (*mEGFPI*) and either pCMV-3Tag-8-Barr (control), pALAF015 (WT), or pALAF030 (dRNase). The retrotransposition efficiency was normalized to the transfection efficiency control (*i.e.*, cells co-transfected with cepB-gfp-L1.3RT(−) intronless and either pCMV-3Tag-8-Barr (control), pALAF015 (WT), or pALAF030 (dRNase)). X-axis, name of the plasmid co-transfected with cepB-gfp-L1.3 (*mEGFPI*). Y-axis, relative retrotransposition efficiency relative to the cepB-gfp-L1.3 (*mEGFPI*) + pCMV-3Tag-8-Barr control. Pairwise comparisons relative to the control: *p* = 9.5 × 10^−5^*** (WT); 0.0073** (dRNase). **(d)** *L1 retrotransposition efficiency in the presence of the D1346N/D1354N/D1355N (dRNase) mutant in HEK293T cells*. Experiments were conducted as summarized in panel (c). Pairwise comparisons relative to the cepB-gfp-L1.3 (*mEGFPI*) + pCMV-3Tag-8-Barr control: *p* = 9.4 × 10^−10^*** (WT HELZ2), 4.1 × 10^−8^*** (dRNase). Values represent the mean ± SEM of three independent biological replicates. The *p*-values were calculated using a one-way ANOVA followed by Bonferroni-Holm post-hoc tests. ns: not significant; *** *p<*0.001.

**Supplementary Figure. 6 (supporting Figs. 6b, 6d, 6e, and 6f):**
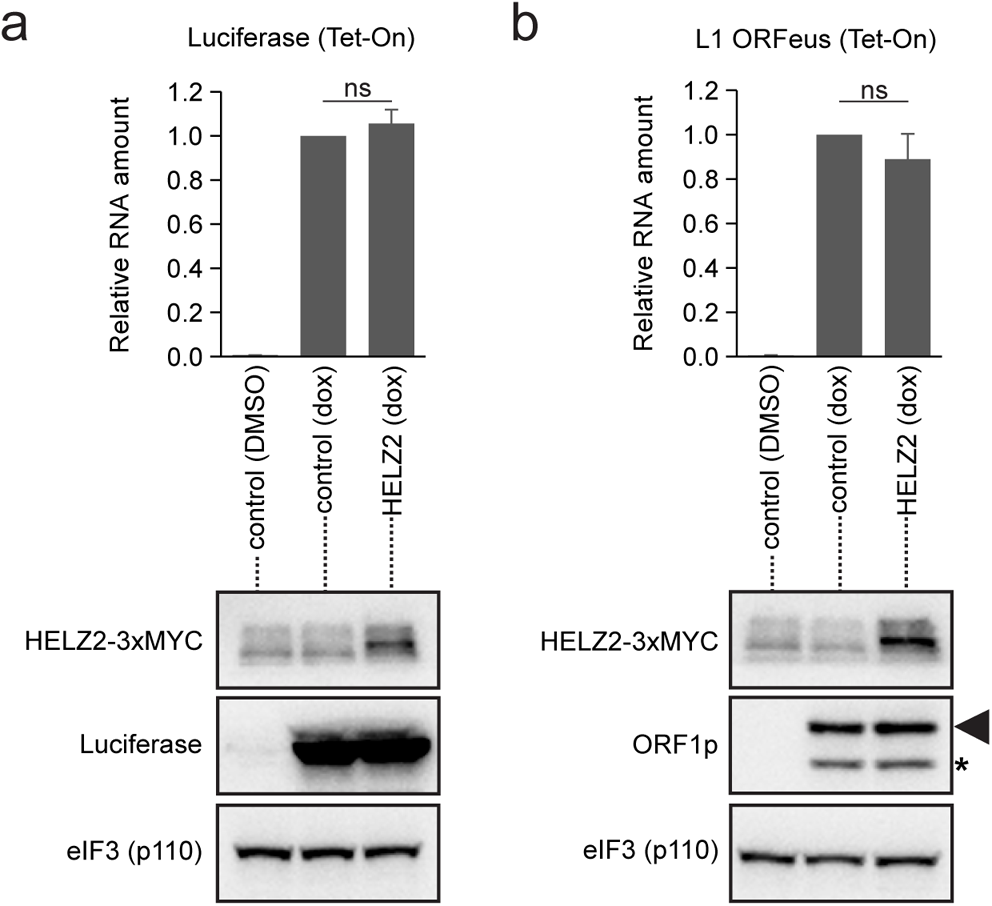
The L1 5’UTR is required for the HELZ2-mediated reduction in L1 RNA steady state levels. **(a & b)** *The effect of HELZ2 on doxycycline inducible (Tet-On) luciferase (panel [a]) or human L1 ORFeus (panel [b]) expression.* HeLa-JVM cells expressing inducible firefly luciferase (pSBtet-RN) or human L1 ORFeus (pDA093) were treated with vehicle (DMSO) or doxycycline (dox) and then transfected with either pCMV-3Tag-8-Barr (control) or pALAF015 (HELZ2). Cells were collected 48 hours post-transfection. Top: Luciferase and L1 levels were quantified using RT-qPCR (primer set: Luciferase and L1 [SV40], respectively) and normalized to *GAPDH* RNA levels (primer set: GAPDH). X-axis, construct name and whether cells were treated with vehicle (DMSO) or doxycycline (dox). Y-axis, RNA levels normalized to the inducible firefly luciferase (pSBtet-RN) or human L1 ORFeus (pDA093) + pCMV-3Tag-8-Barr control. Bottom: western blot analyses. An anti-MYC antibody was used to detect HELZ2, an anti-luciferase antibody was used to detect luciferase, and an anti-ORF1p antibody was used to detect ORF1p. Black arrowhead (middle right blot), the expected ORF1p band; asterisk (middle right blot), unexpected lower molecular weight ORF1p band. The eIF3 subunit (p110) served as a loading control. Values in the graphs represent the mean ± SEM of three independent biological replicates. The *p*-values were calculated using a one-way ANOVA followed by Bonferroni-Holm post-hoc tests: *p* = 0.32^ns^ (Luciferase); and 0.28^ns^ (L1); ns: not significant.

